# Inflammation causes insulin resistance via interferon regulatory factor 3 (IRF3)-mediated reduction in FAHFA levels

**DOI:** 10.1101/2023.08.08.552481

**Authors:** Shuai Yan, Anna Santoro, Micah J. Niphakis, Antonio M. Pinto, Christopher L. Jacobs, Rasheed Ahmad, Radu M. Suciu, Bryan R. Fonslow, Rachel B. Herbst-Graham, Nhi Ngo, Cassandra L. Henry, Dylan M. Herbst, Alan Saghatelian, Barbara B. Kahn, Evan D. Rosen

## Abstract

Obesity-induced inflammation causes metabolic dysfunction, but the mechanisms remain elusive. Here we show that the innate immune transcription factor interferon regulatory factor (IRF3) adversely affects glucose homeostasis through induction of the endogenous FAHFA hydrolase androgen induced gene 1 (AIG1) in adipocytes. Adipocyte-specific knockout of IRF3 protects mice against high-fat diet-induced insulin resistance, whereas overexpression of IRF3 or AIG1 in adipocytes promotes insulin resistance on a high-fat diet. Furthermore, pharmacological inhibition of AIG1 reversed obesity-induced insulin resistance and restored glucose homeostasis in the setting of adipocyte IRF3 overexpression. We, therefore, identify the adipocyte IRF3/AIG1 axis as a crucial link between obesity-induced inflammation and insulin resistance and suggest an approach for limiting the metabolic dysfunction accompanying obesity.

## Main

In the setting of persistent caloric excess, pathological white adipose tissue (WAT) expansion is accompanied by a state of chronic inflammation, sometimes termed “meta-inflammation”, leading to insulin resistance and, potentially, type 2 diabetes^1^. WAT inflammation is associated with recruitment^2, 3^ and proliferation^4^ of macrophages and other immune cells, which secrete a variety of proinflammatory cytokines^5^. These cytokines, combined with other inflammatory stimuli that increase in obesity (e.g., LPS from the gut and cell-free DNA from dying adipocytes)^6, 7^, cause insulin resistance and related metabolic dysfunction.

Despite clear data demonstrating the role of inflammation in insulin resistance, the mechanism by which this occurs is still unclear. Most attention has focused on the acute effects of inflammation on proximal insulin signaling, despite evidence suggesting that the time course over which cytokine signals work (minutes to a few hours) is not concordant with the time course over which insulin action is affected (several hours to days)^8^. An alternative explanation is that inflammatory factors activate critical transcriptional pathways that alter the receptiveness of WAT and other tissues to insulin. This notion is bolstered by observations that several transcription factors (e.g., PPARγ) are known to alter insulin sensitivity, an effect shared by drugs or genetic manipulations that affect chromatin structure^9–11^.

It is common to invoke increased activity of nuclear factor κB (NF-κB) in the setting of inflammation. NF-κB is sequestered in the cytosol bound to the inhibitor IκB protein until phosphorylation of IκB by IKKβ releases it to enter the nucleus^12^. Manipulations of IKKβ show the predicted effects on insulin action, especially in the liver^13^. This effect, however, is likely to be independent of NF-κB, given that (a) IKKβ also phosphorylates insulin signaling proteins directly^14^, (b) overexpression of the active p65 subunit of NF-κB in adipocytes maintains insulin sensitivity in both age-related and diet-induced obesity in mice, despite proinflammatory gene expression in adipose tissue^15, 16^, and (c) mice lacking the regulatory p50 subunit have increased NF-κB activity and yet display resistance to diet-induced obesity and preserved insulin action^17, 18^. Taken together, these studies indicate that transcriptional pathways other than NF-κB must mediate the effect of inflammation on insulin resistance. These observations also suggest that the classic proinflammatory cytokine portfolio induced by NF-κB and similar transcription factors may not be driving forces in insulin resistance, as is commonly assumed.

Another class of transcription factors with major roles in both innate and adaptive immunity are the interferon regulatory factors (IRFs), comprised of 9 members (IRF1– IRF9). IRFs have been implicated in a wide range of immune functions, including lymphopoiesis, macrophage differentiation, and antiviral defense^19^. We have previously determined the role of several IRFs in adipogenesis^20^ and further demonstrated that IRF4 is an important regulator of adipocyte lipolysis^21^ and thermogenesis^22^. The family member most associated with transducing proinflammatory signals is IRF3^23^. IRF3 expression is induced in the adipose tissue of people and rodents with obesity^24^, and mice lacking IRF3 globally are resistant to diet-induced obesity, an effect recapitulated in mice lacking IRF3 specifically in adipocytes. Conversely, expression of a constitutively active form of IRF3 in adipocytes promotes weight gain^25^. This effect on adiposity is mediated by direct activation of *Isg15* gene expression and protein ISGylation in adipocytes, which results in reduced glycolysis and lactate production and strongly dampens adipose thermogenesis^25^. In addition to repressing thermogenesis, IRF3 in adipocytes may also exert weight- and ISGylation-independent effects on insulin sensitivity and glucose homeostasis, although this was not proven definitively, and no downstream mechanism has been identified^25^. This is in contrast to liver, where IRF3 was shown to impact hepatic insulin action through the direct induction of the protein phosphatase 2A (PP2A) subunit *Ppp2r1b*^26^.

Here we find that inflammatory stimuli modulate insulin action in adipocytes in a cell autonomous and IRF3-dependent manner. High-fat fed adipocyte-specific *Irf3* deficient (FI3KO) mice studied at thermoneutrality exhibit improved insulin and glucose tolerance, while adipocyte-specific overexpression of *Irf3* (FI3OE) promotes glucose intolerance and insulin resistance on HFD, also in a weight-independent manner. Mechanistically, we identify AIG1, an endogenous hydrolase of fatty acyl esters of hydroxy fatty acids (FAHFAs), as a downstream target of IRF3. Overexpression of AIG1 in adipose tissue impairs insulin and glucose tolerance in FI3KO mice. In addition, a novel small molecule inhibitor of AIG1 raises FAHFA levels and attenuates HFD-induced insulin resistance and glucose intolerance in FI3OE mice. These findings uncover a defined and direct mechanism by which inflammation impairs adipose insulin sensitivity and glucose homeostasis in obesity.

## Results

### Activation of toll-like receptors 3 and 4 provokes insulin resistance in cultured adipocytes in an IRF3-dependent manner

TLR4 activation has been shown to reduce insulin-stimulated glucose uptake in adipocytes^27^. We first sought to recapitulate this effect in cultured adipocytes derived from mouse stromal vascular fraction (SVF). Treatment with LPS for 2 days led to a dose-dependent decrease in insulin-stimulated glucose uptake (**Fig. 1a**). LPS treatment also increased phosphorylation of murine IRF3 at serine 388 (**Fig. 1b**), which is known to promote nuclear translocation, dimerization, and binding to DNA^28^.

**Fig. 1.**
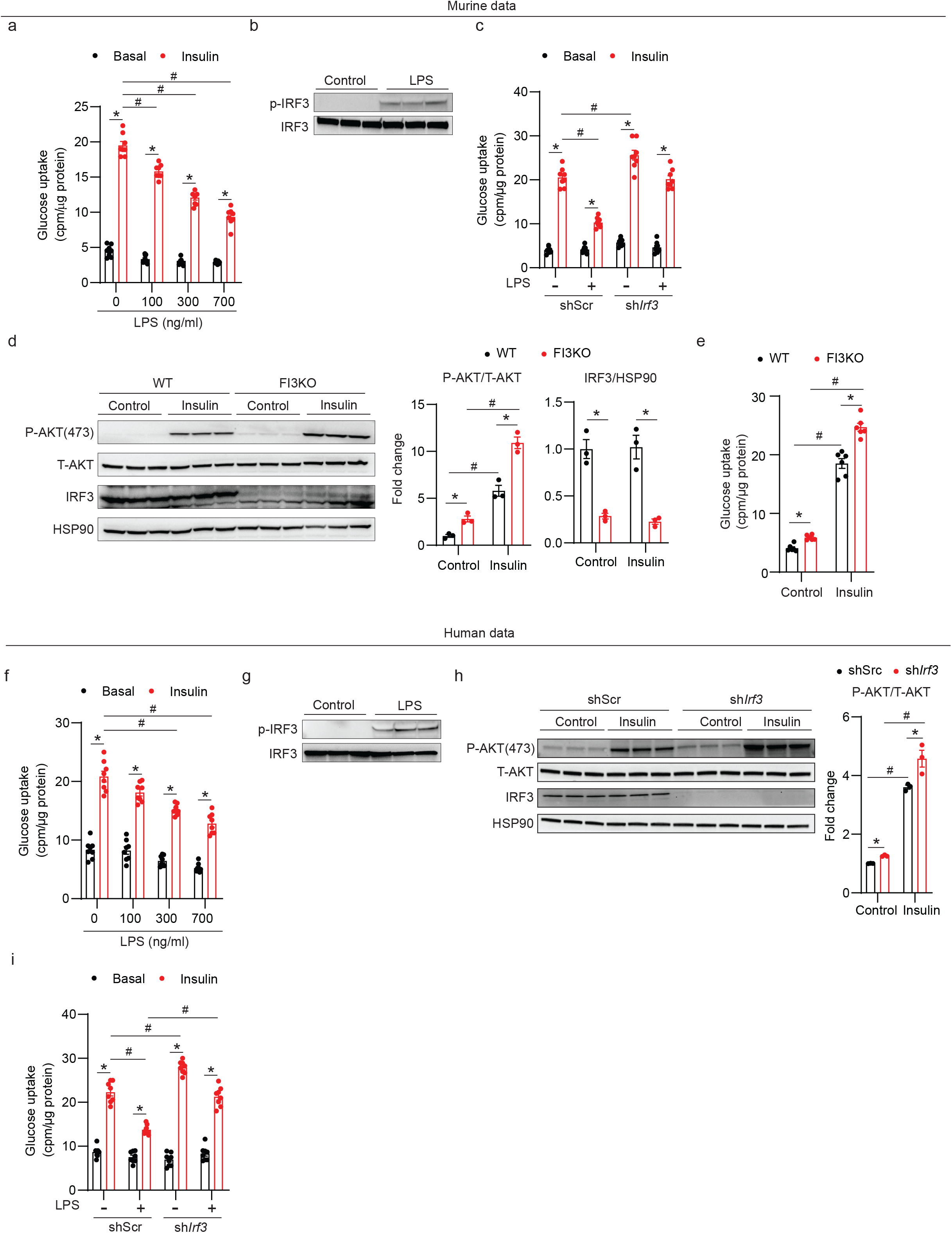
IRF3 mediates insulin resistance in response to TLR4 ligands in cultured adipocytes. **a,** Glucose uptake in mouse adipocytes after treatment with varying doses of LPS for 2 days (n=8). **b,** Western blot showing phosphorylation of murine IRF3 (S388) in mouse adipocytes after 30 mins of LPS (700 ng/ml) treatment. **c,** Glucose uptake in mouse adipocytes transduced with lentivirus expressing shRNA against *Irf3* or shScr control hairpin in the absence or presence of LPS (700 ng/ml) (n=8). d, Western blot of pAKT (S473) and IRF3, **e,** Glucose uptake (n=6) in WT and FI3KO SVF-derived adipocytes treated with control or 100nM insulin. **f,** Glucose uptake in human SGBS adipocytes after treatment with varying doses of LPS for 2 days (n=8). **g,** Western blot showing phosphorylation of human IRF3 (S396) in human SGBS adipocytes after 30 mins of LPS (700 ng/ml) treatment. **h,** Western blot of pAKT (S473) and IRF3 in human SGBS adipocytes transduced with lentivirus expressing shRNA against *Irf3* or a scrambled control hairpin (shScr). **i,** Glucose uptake in human SGBS adipocytes transduced with lentivirus expressing shRNA against *Irf3* or shScr control hairpin in the absence or presence of LPS (700 ng/ml) (n=8). Statistical significance was assessed by *three-way* ANOVA (**c** and **i**) or *two-way* ANOVA (**a**, **d**, **e**, **f**, and **h**). Data are expressed as mean ± SEM. *P < 0.05 vs WT, ^#^P < 0.05 vs. control.

To determine whether IRF3 is required for LPS-induced insulin resistance, we knocked down IRF3 in mature adipocytes using a doxycycline-inducible lentivirus. Knocking down IRF3 abolished the suppressive effect of LPS treatment on glucose uptake (**Fig. 1c**). We next isolated primary mouse inguinal SVF cells from *Irf3^flox/flox^*mice for *in vitro* differentiation into mature adipocytes, and then infected these cells with AAV-GFP (WT) or AAV-GFP-Cre (FI3KO). In mature adipocytes, IRF3 deletion led to a robust decrease in the expression of IRF3 target genes, including *Ccl5*, *Ifit1,* and *Isg15* (**Supplementary Fig. 1a**). Importantly, there was no significant difference in differentiation state between WT and FI3KO cells, as determined by the expression of mature adipocyte marker genes (**Supplementary Fig. 1b**). We observed higher p-AKT levels in FI3KO adipocytes compared with WT cells even in the absence of insulin, supporting the idea that endogenous adipocyte IRF3 represses basal Akt phosphorylation (**Fig. 1d**). Consistent with the signaling data, ablation of IRF3 enabled increased insulin-induced glucose uptake; a much smaller effect of IRF3 on glucose uptake was also observed in the absence of insulin (**Fig. 1e**). Finally, FI3KO adipocytes displayed greater sensitivity to the anti-lipolytic effects of insulin (**Supplementary Fig. 1c**).

To confirm these findings in human adipocytes, we used the Simpson-Golabi-Behmel syndrome (SGBS) preadipocyte cell strain^29^. As seen in mouse adipocytes, LPS treatment of human adipocytes led to a dose-dependent decrease in insulin-stimulated glucose uptake and an increase in IRF3 phosphorylation (**Fig. 1f, g)**. Knockdown of IRF3 in SGBS cells abolished the suppressive effect of LPS treatment on insulin sensitivity, as measured by p-Akt and glucose uptake (**Fig. 1h, i**).

IRF3 can also be activated through TLR3 signaling [e.g., polyinosinic-polycytidylic acid (poly I:C) treatment]^30^. We therefore determined the effect of the TLR3/IRF3 signaling axis in regulating glucose uptake in mouse SVF-derived adipocytes and human SGBS cells. Poly I:C treatment robustly decreased insulin-stimulated glucose uptake and increased phosphorylation of IRF3 in mouse SVF-derived adipocytes (**Supplementary Fig. 1d, e**). shRNA-mediated reduction of IRF3 fully abrogated poly I:C–induced insulin resistance in these cells (**Supplementary Fig. 1f**). Similar data were obtained from human adipocytes treated with poly I:C (**Supplementary Fig. 1g-i**).

### Constitutively active IRF3 induces insulin resistance in mouse and human adipocytes

We have shown that the double-mutant murine *Irf3* allele (S388D/S390D; hereafter designated as IRF3-2D) is constitutively active *in vitro*^25^. Concordant with the loss-of-function studies just described, overexpression of IRF3-2D in mouse SVF-derived adipocytes led to a significant decrease in insulin-stimulated glucose uptake and p-Akt in the absence of upstream activators (**Fig. 2a**). We also isolated mouse inguinal SVF cells from Irf3-2D mice, differentiated them into mature adipocytes, and infected these cells with AAV-GFP (WT) or AAV-GFP-Cre (FI3OE). IRF3 overexpression in mature adipocytes markedly increased the expression of target genes, without affecting adipogenesis (**Supplementary Fig. 2a, b**). FI3OE cells displayed decreased insulin-stimulated AKT phosphorylation and glucose uptake compared with WT cells; insulin was also less able to suppress lipolysis in these cells (**Fig. 2b, c** and **Supplementary Fig. 2c**). Human SGBS cells behaved similarly when IRF3-2D was expressed, showing reduced insulin-stimulated p-AKT and glucose uptake (**Fig. 2d, e)** in SGBS cells. Taken together with the loss-of-function studies, these data strongly suggest that adipocyte IRF3 is both necessary and sufficient to promote insulin resistance in adipocytes, and that it does so in a cell-autonomous and species-independent manner.

**Fig. 2.**
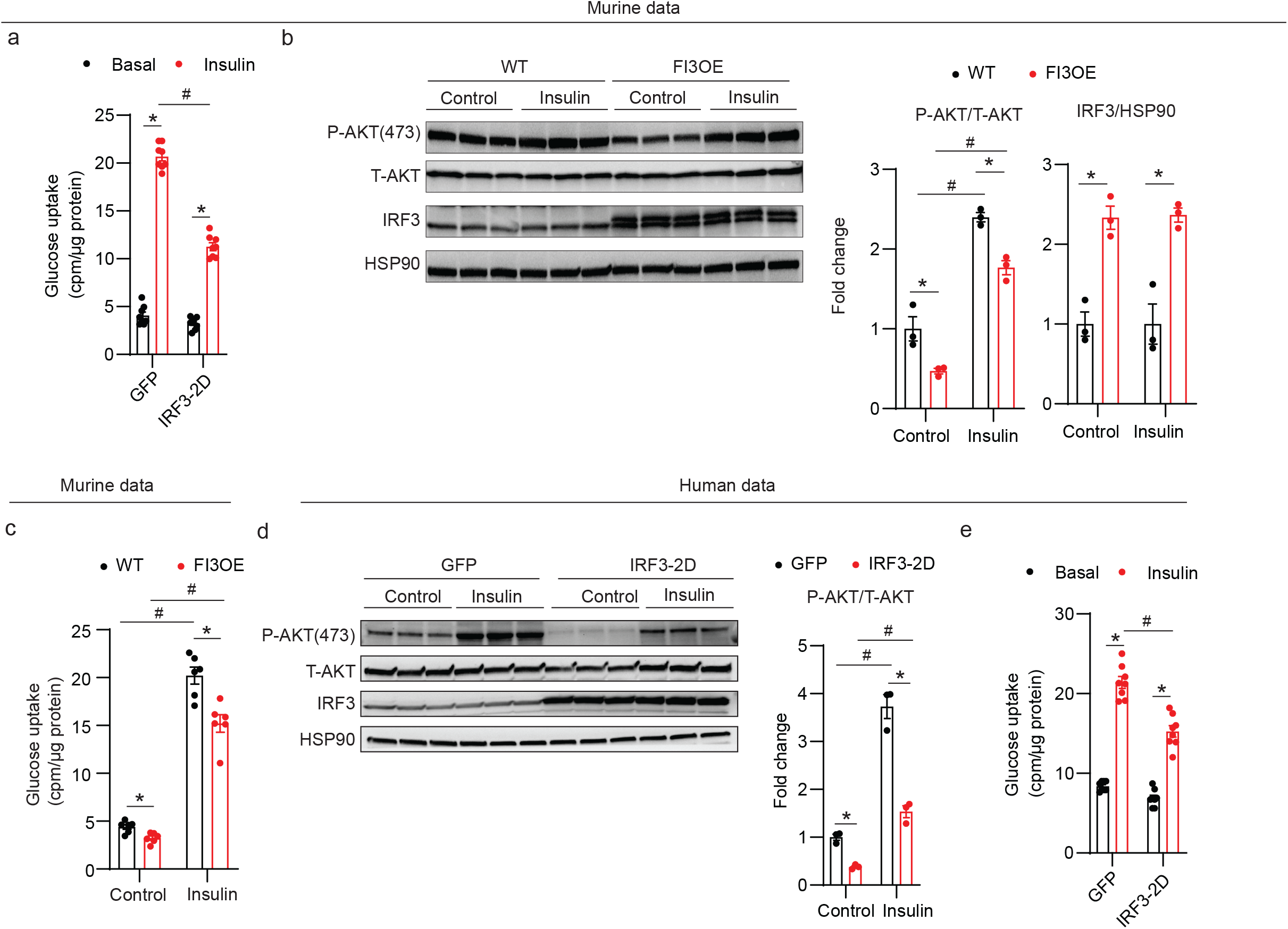
IRF3 promotes insulin resistance in cultured mouse and human adipocytes. **a,** Glucose uptake in mouse adipocytes expressing GFP or constitutively active IRF3-2D mutant (n=8). **b,** Western blot of pAKT (S473) and IRF3 in WT and FI3OE SVF-derived adipocytes treated with control or 100nM insulin. **c,** Glucose uptake (n=6) in WT and FI3OE SVF-derived adipocytes treated with vehicle or 100nM insulin. **d,** Western blot of pAKT (S473) and IRF3 in human SGBS adipocytes transduced with lentivirus expressing GFP or IRF3-2D. **e,** Glucose uptake in human SGBS adipocytes expressing GFP or IRF3-2D mutant (n=8). Statistical significance was assessed by *two-way* ANOVA. Data are expressed as mean ± SEM. *P < 0.05 vs WT, ^#^P < 0.05 vs. control.

### Adipocyte-specific knockout of *Irf3* attenuates HFD-induced insulin resistance in mice at thermoneutrality

In our previous work, we demonstrated that fat-specific IRF3 knockout (FI3KO) mice have increased adipose thermogenesis and are protected from diet-induced obesity^25^, which confounds the ability to determine whether changes in insulin sensitivity are dependent or independent of changes in adiposity. To avoid this issue, we housed mice under thermoneutral (TN) conditions (30°C) at which there is no requirement for adaptive thermogenesis to maintain body temperature (**Supplementary Fig. 3a**)^31^. After 16 weeks of HFD feeding, weight gain, adiposity, and food intake were not different between male FI3KO and WT mice (**Supplementary Fig. 3b-d**). However, despite the absence of protection from diet-induced obesity, FI3KO mice exhibited markedly improved insulin sensitivity and glucose tolerance (**Fig. 3a-c**). Consistent with the improved glycemia, FI3KO mice exhibited significantly increased insulin-stimulated AKT phosphorylation within epidydimal white adipose tissue (eWAT), inguinal WAT (iWAT), skeletal muscle, and liver, but not brown adipose tissue (BAT) (**Fig. 3d**). To understand the tissue-specific basis of the insulin sensitivity phenotype in FI3KO mice at a more quantitative level, we performed insulin-stimulated glucose uptake of ^14^C-2-deoxyglucose *in vivo* in HFD-fed mice. Glucose uptake by BAT, iWAT, eWAT, quadriceps muscle, and heart was significantly increased in FI3KO mice compared to WT controls, indicating that the improved insulin sensitivity in FI3KO mice may result from greater glucose uptake into a wide range of insulin-sensitive tissues (**Fig. 3e**). Taken together, these results indicate that IRF3 deficiency in adipocytes is sufficient to attenuate obesity-induced insulin resistance at thermoneutrality.

**Fig. 3.**
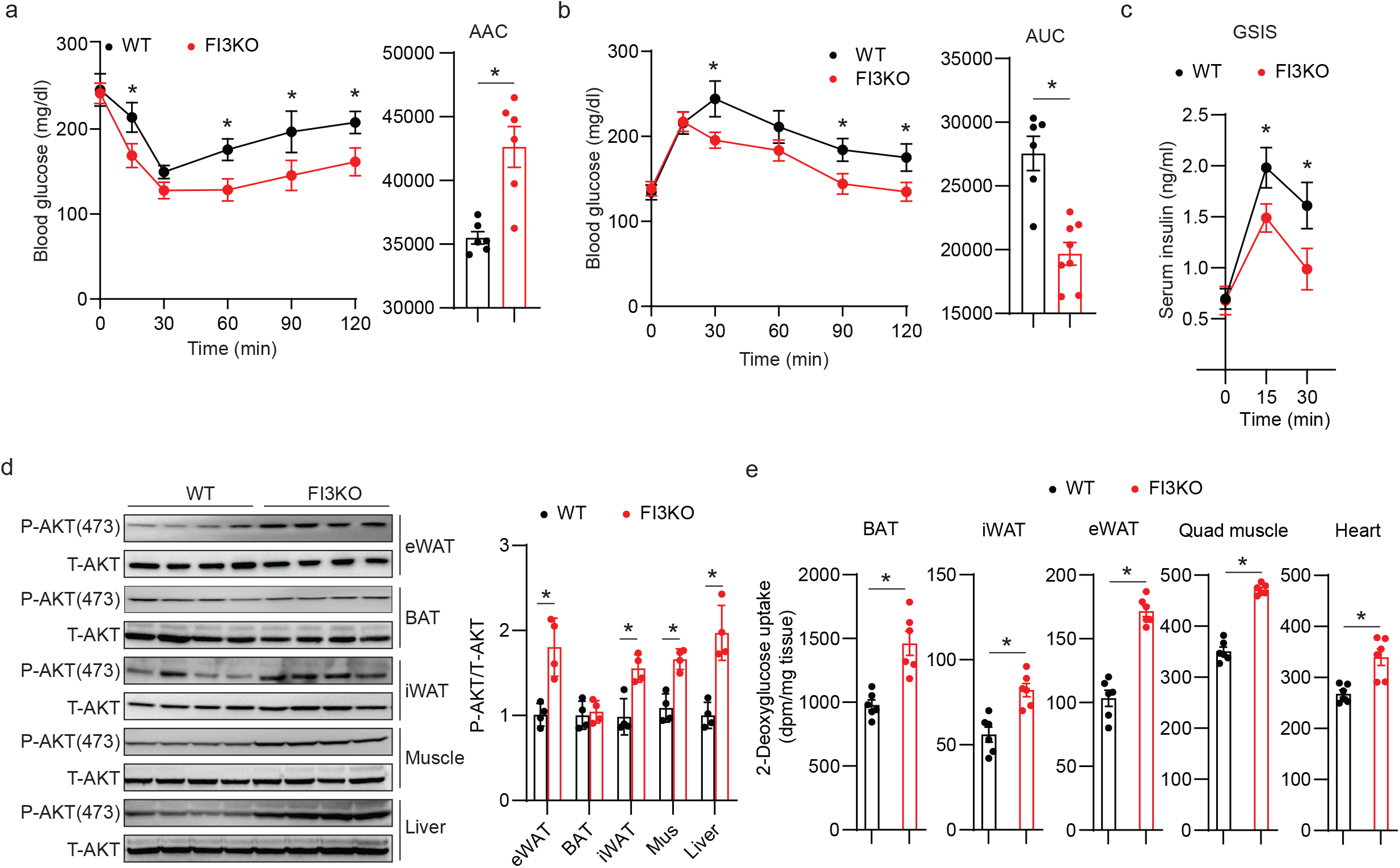
Adipocyte-specific knockout of *Irf3* attenuates HFD-induced insulin resistance in mice at thermoneutrality. Metabolic analysis of male WT and FI3KO mice (n=10) after 16 weeks of HFD feeding at thermoneutrality, including insulin tolerance test (**a**), glucose tolerance test (**b**), *ad lib* fed serum insulin levels (**c**), immunoblotting for P-AKT (S473) protein levels in eWAT, BAT, iWAT, muscle, and liver (**d**), in *vivo* 2-deoxyglucose uptake in BAT, iWAT, eWAT, quadriceps muscle and heart (**e**). Statistical significance was assessed by *two-way* ANOVA (**a**-**c**) or two-tailed Student’s *t-test* (**d** and **e**). Data in all panels are expressed as mean ± SEM. *P < 0.05 vs WT.

### Adipocyte-specific overexpression of *Irf3* promotes HFD-induced insulin resistance in mice at thermoneutrality

To further investigate the role of adipocyte IRF3 in insulin sensitivity, we used a line of knock-in mice (FI3OE) in which IRF3 is constitutively activated in adipocytes without the need for an external stimulus like LPS or poly I:C. FI3OE mice display enhanced weight gain when housed at 23°C^25^. After 16 weeks of HFD feeding at thermoneutrality, however, weight gain, adiposity, and food intake were comparable between FI3OE and WT mice (**Supplementary Fig. 3e-g**). Despite weighing the same as control mice, FI3OE mice exhibited markedly impaired insulin sensitivity and glucose tolerance (**Fig. 4a-c**). FI3OE mice exhibited a significant decrease in insulin-stimulated AKT phosphorylation in eWAT, BAT, iWAT, skeletal muscle, and liver (**Fig. 4d**). Consistent with this, glucose uptake by BAT, iWAT, eWAT, quadriceps muscle, and heart was significantly decreased in FI3OE mice compared to WT controls (**Fig. 4e**).

**Fig. 4.**
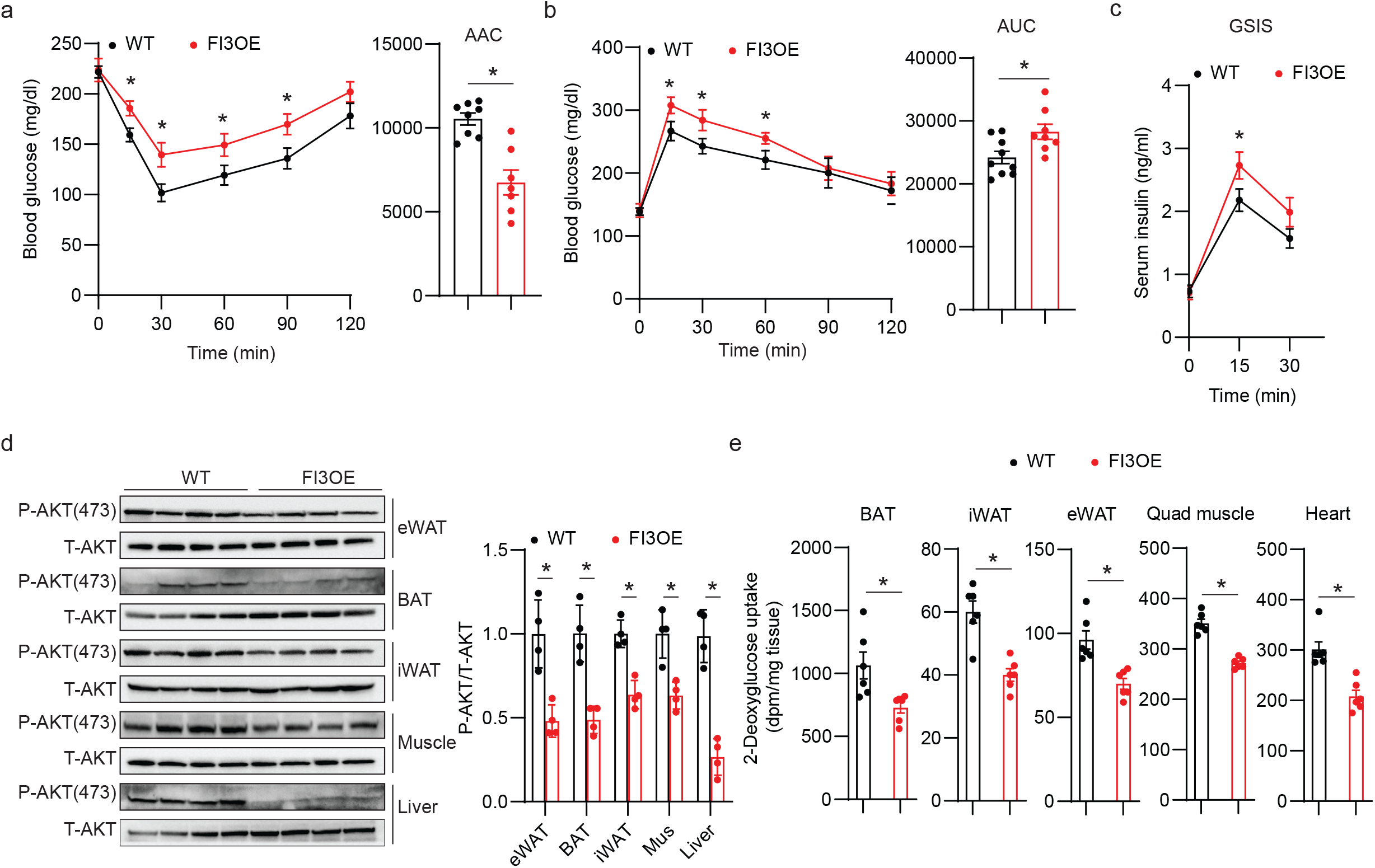
Adipocyte IRF3 promotes HFD-induced insulin resistance at thermoneutrality. (**a**-**e**) Metabolic analysis of male WT and FI3OE mice (n=10) after 16 weeks of HFD feeding at thermoneutrality, including insulin tolerance test (**a**), glucose tolerance test (**b**), serum insulin levels (**c**), immunoblotting for pAKT (S473) protein levels in eWAT, BAT, iWAT, muscle and liver (**d**), *in vivo* 2-deoxyglucose uptake in BAT, iWAT, eWAT, quadriceps muscle, and heart (**e**). Statistical significance was assessed by *two-way* ANOVA (**a**-**c**) or two-tailed Student’s *t-test* (**d** and **e**). Data in all panels are expressed as mean ± SEM. *P < 0.05 vs WT.

### IRF3 drives AIG1 expression, leading to reductions in intracellular FAHFA levels

The finding that manipulating IRF3 levels in adipocytes affects insulin sensitivity in other tissues suggested that IRF3 might alter adipokine secretion or action. To address this possibility, we co-cultured WT SVF-derived adipocytes with conditioned medium from either WT SVF-derived adipocytes (WT-CM), FI3KO SVF-derived adipocytes (FI3KO-CM), or FI3KO SVF-derived adipocytes treated with proteinase K (FI3KO-CM+PK). Surprisingly, both FI3KO-CM and FI3KO-CM+PK markedly increased p-AKT in WT adipocytes compared with WT-CM, indicating that a protease-insensitive substance(s) from FI3KO adipocytes can enhance insulin signaling (**Fig. 5a**).

**Fig. 5.**
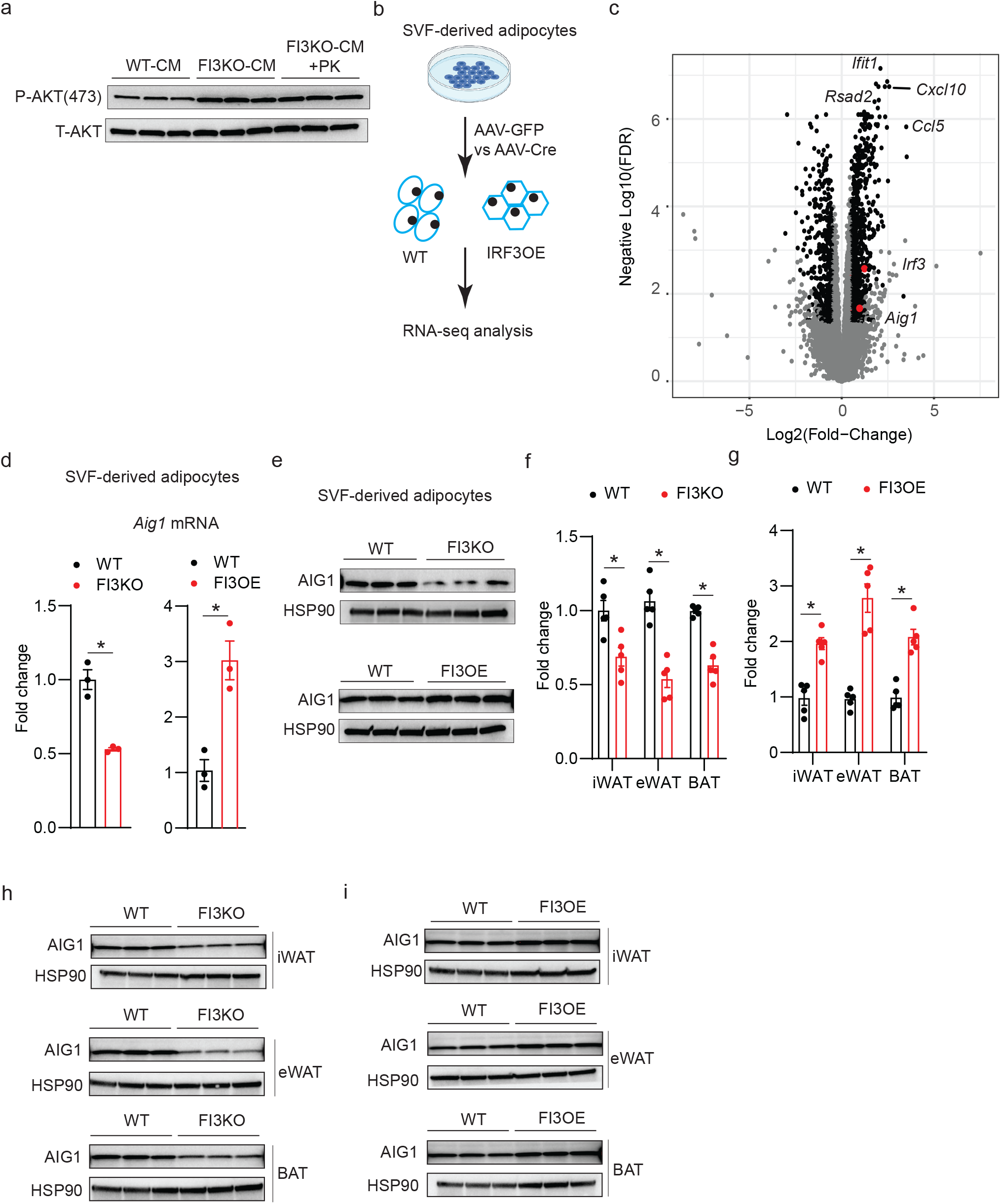
IRF3 increases the transcription of AIG1 in adipocytes. **a,** Western blot of pAKT (S473) in WT SVF-derived adipocytes pre-treated with indicated conditioned medium plus insulin (100nM) for 30 mins. **b,** Scheme showing the experimental paradigm used to identify IRF3 target genes by RNA-sequencing. **c,** Volcano plot of RNA-seq dataset from **b**. Genes that meet both significance and abundance are shown in black. *Irf3* and *Aig1* are marked in red. **d** and **e,** mRNA levels of *Aig1* (n=3) (**d**) and western blot (**e**) of AIG1 in WT and FI3KO, FI3OE SVF-derived adipocytes. **f** and **g,** mRNA levels of *Aig1* in iWAT, eWAT, and BAT from WT, FI3KO (**f**) and FI3OE mice (**g**) (n=5). **h,** Western blot of AIG1 in iWAT, eWAT, and BAT of WT and FI3KO mice (n=3). **i,** Western blot of AIG1 in iWAT, eWAT, and BAT of WT and FI3OE mice (n=3). Statistical significance was assessed by two-tailed Student’s *t-test*. Data in all panels are expressed as mean ± SEM. *P < 0.05 vs WT.

IRF3 is a transcription factor, and so we sought to identify gene targets that could be credibly linked to changes in levels of insulin-sensitizing metabolites or lipids. To that end, we defined the transcriptional profiles of WT and FI3OE SVF-derived adipocytes (**Fig. 5b** and **Extended Data Table 1**). One significantly affected gene was androgen-induced gene 1 (*Aig1*) (**Fig. 5c**), which encodes a hydrolase that degrades members of the fatty acyl ester of hydroxy fatty acid (FAHFA) family of lipids produced by adipocytes, which have been implicated as important endogenous insulin sensitizers^32–34^. qPCR and western blotting of SVF-derived adipocytes from FI3KO and FI3OE mice confirmed altered *Aig1* expression (**Fig. 5d, e**). Furthermore, *Aig1* was down-regulated in iWAT, eWAT, and BAT from FI3KO mice, and up-regulated in FI3OE mice (**Fig. 5f-i**). *Aig1* expression in other tissues, such as skeletal muscle, liver, and heart, was not altered in FI3KO and FI3OE mice (**Supplementary Fig. 4a, b**). Expression of another known FAHFA hydrolase, androgen-dependent TFPI-regulating protein (ADTRP), was unchanged in the tissues of FI3KO and FI3OE mice (**Supplementary Fig. 4c, d**).

We next used targeted lipidomics to quantify isomers from multiple FAHFA families, many of which are known to be important in insulin action and glucose homeostasis. We found that SVF-derived FI3KO adipocytes show elevated levels of several FAHFA species, including oleic acid-hydroxy stearic acids (OAHSAs), palmitic acid-hydroxy stearic acids (PAHSAs), and palmitoleic acid-hydroxy stearic acids (POHSAs) (**Fig. 6a** and **Supplementary Fig. 5a**). Conversely, reduced FAHFA levels of these same families were observed in FI3OE adipocytes (**Fig. 6b** and **Supplementary Fig. 5b**). Underscoring the physiological relevance of these in vitro results, we found that FAHFA levels were also markedly elevated in vivo in whole eWAT of FI3KO mice (**Fig. 6c** and **Supplementary Fig. 5c**), and decreased in the eWAT of FI3OE mice (**Fig. 6d** and **Supplementary Fig. 5d**).

**Fig. 6.**
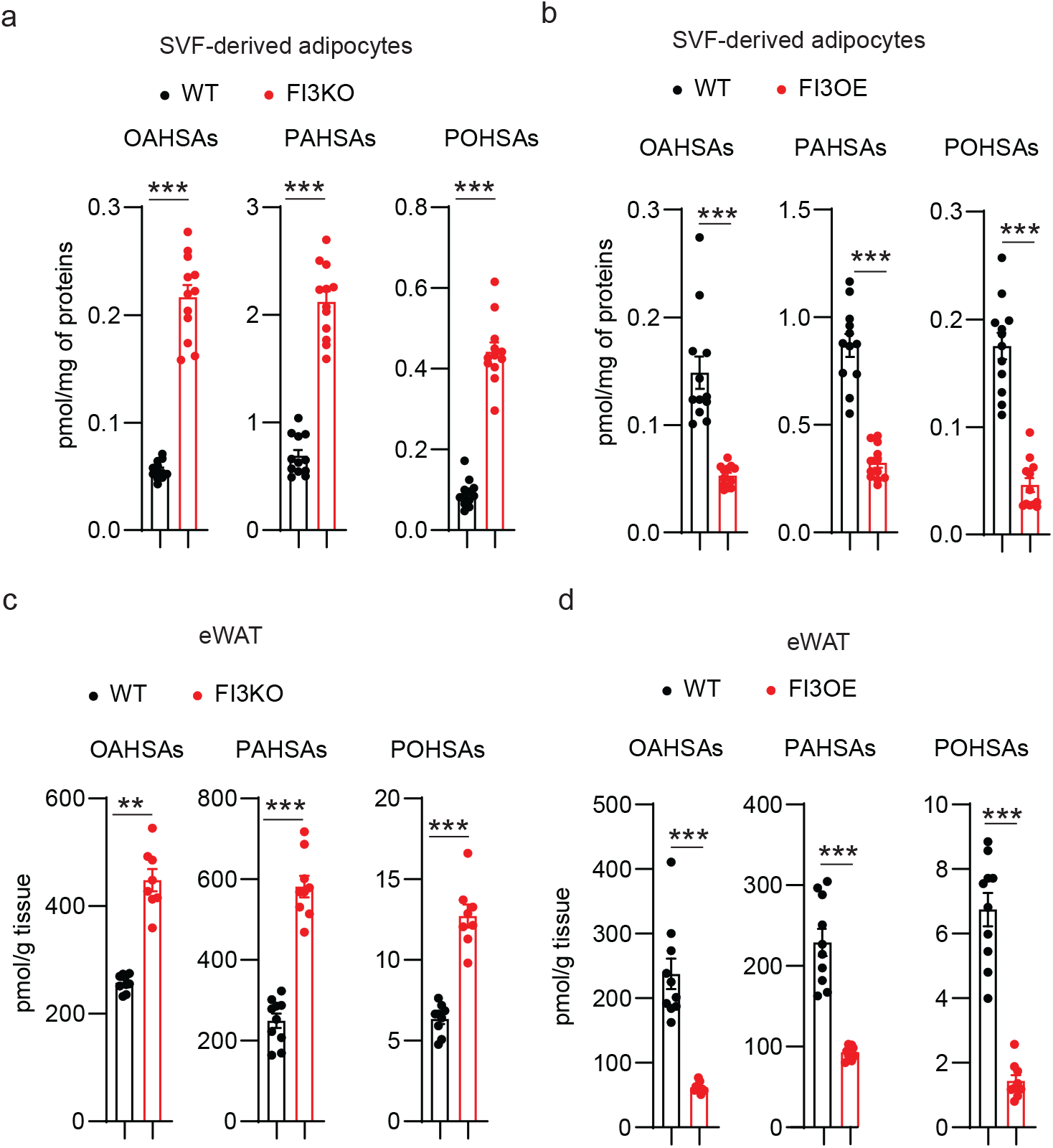
IRF3 decreases intracellular FAHFAs levels. **a**, Cellular FAHFA levels of WT and FI3KO and **b**, WT and FI3OE SVF-derived adipocytes (n=12). **c**, Tissue FAHFA levels of eWAT from WT and FI3KO and **d**, WT and FI3OE mice (n=10). Statistical significance was assessed by two-tailed Student’s t-test. Data in all panels are expressed as mean ± SEM. **P < 0.01 vs WT, ***P < 0.001 vs WT.

### AIG1 is responsible for IRF3-mediated impaired insulin sensitivity

We next tested whether AIG1 is required for IRF3-mediated suppression of insulin sensitivity *in vitro*. As before, FI3KO adipocytes exhibited elevated p-AKT level and insulin-stimulated glucose uptake, which was fully suppressible by WT AIG1 overexpression, but not two different alleles of *Aig1* bearing mutations in the catalytic domain (T43A and H134A) (**Supplementary Fig. 6a, b**)^33^. Conversely, the reduced p-AKT levels and insulin-stimulated glucose uptake observed in FI3OE adipocytes were rescued by knockdown of AIG1 (**Supplementary Fig. 6c, d**). In addition, decreased glucose uptake was observed in FI3OE SVF-derived adipocytes rescued by FAHFAs (**Supplementary Fig. 6e**).

To clarify the role of AIG1 in mediating the metabolic functions of IRF3 *in vivo*, we overexpressed AIG1 via AAV-AIG1 (vs. an AAV-GFP control) in eWAT of WT and FI3KO mice (**Fig. 7a**). AIG1 overexpression in eWAT did not affect the body weight of WT and FI3KO mice, but largely blunted the metabolic improvements of FI3KO mice (**Fig. 7b-e**). AIG1 overexpression in eWAT also reduced the enhanced p-AKT levels of FI3KO mice in all tissues tested (eWAT, iWAT, BAT, muscle, and liver; **Fig. 7f**). These data suggest that the adipocyte IRF3-AIG1 axis promotes obesity-induced insulin resistance in mice.

**Fig. 7.**
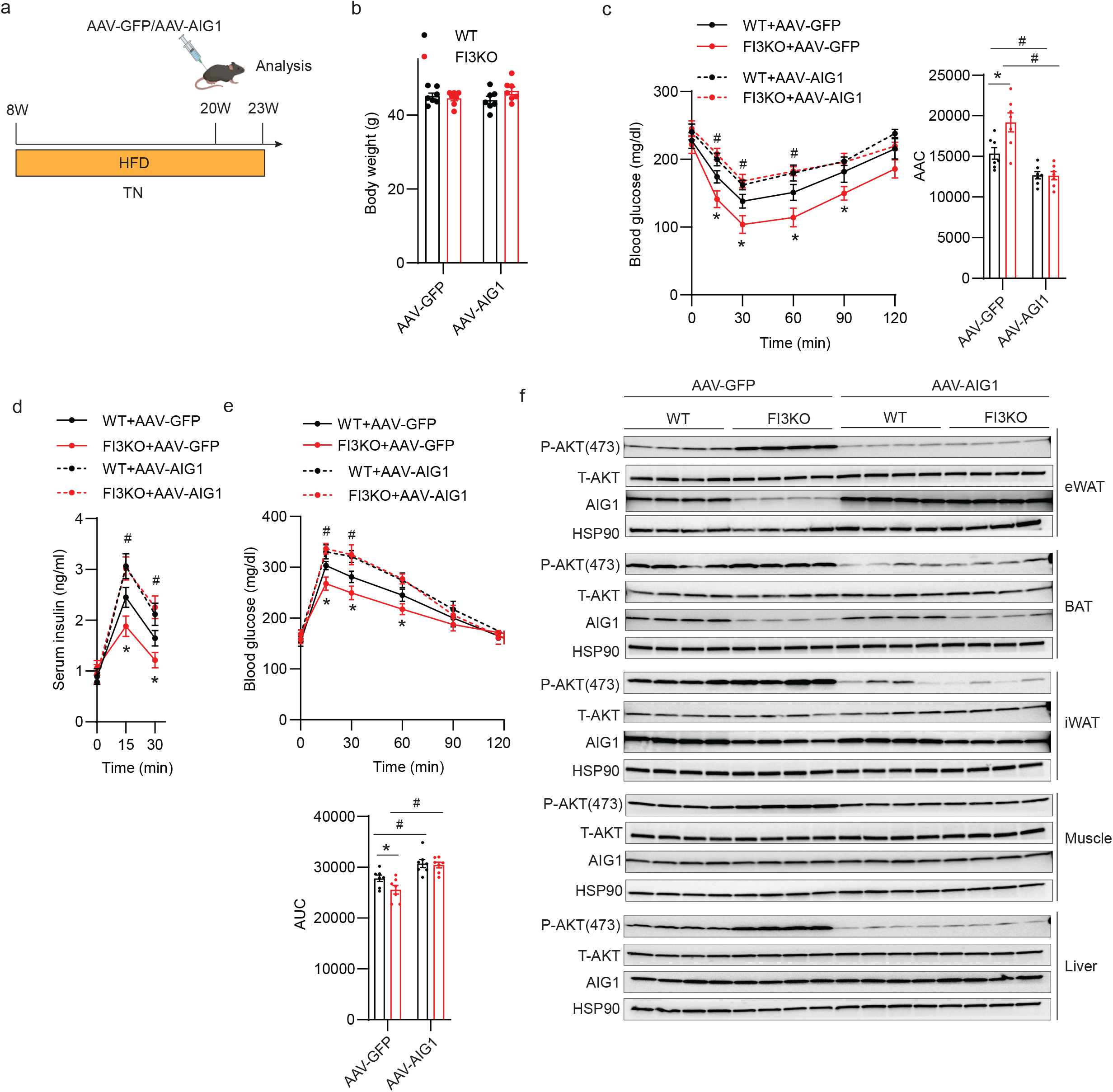
AIG1 is responsible for adipocyte IRF3-mediated suppression of HFD-induced insulin resistance. **a,** Schematic illustrating the strategy for the AAV injection in eWAT of WT and FI3KO mice on HFD at thermoneutrality. Body weight (**b**), insulin tolerance test (**c**), glucose tolerance test (**d**), and serum insulin levels (**e**) of mice described in (**a**) (n=7-8). **f,** Immunoblotting for pAKT (S473) protein levels in mice’s eWAT, BAT, iWAT, muscle, and liver as described in (**a)**. Statistical significance was assessed by *two-way* ANOVA. Data in all panels are expressed as mean ± SEM. *P < 0.05 vs WT, ^#^P < 0.05 vs. GFP.

### Pharmacologically inhibiting AIG1 attenuates HFD-induced insulin resistance in FI3OE mice

To explore the therapeutic potential of inhibiting AIG1, we next sought to develop a selective, *in vivo*-active, AIG1 inhibitor. While a potent, dual AIG1/ADTRP inhibitor (ABD-110207) was previously reported enabling simultaneous *in vivo* evaluation of these FAHFA hydrolases^34^, an in vivo-active AIG1 inhibitor lacking ADTRP activity has yet to be described. Both AIG1 and ADTRP feature an active-site threonine residue which enables both inhibitor screening by activity-based protein profiling (ABPP) using fluorophosphonate (FP) probes and inhibitor design through the application of tailored reactive groups that covalently modify active site nucleophiles^33, 34^. Screening a collection of diverse serine hydrolase-directed compounds by ABPP in mouse brain proteomes, we identified ABD-110000 (called ABD-110 herein) as a potential AIG1-selective tool compound (**Supplementary Fig. 7a**). Given the structural similarity of ABD-110 to other covalent serine hydrolase inhibitors^35, 36^, we believe this compound inactivates AIG1 through carbamylation of the active-site threonine residue **(Fig. 8a**). Gel-based ABPP profiling of ABD-110 in mouse brain revealed high potency against mAIG1 (IC_50_ = 1.6 nM) and selectivity across other serine hydrolases visible by gel imaging; spiking in ADTRP to these samples further revealed the specificity of the inhibitor (**Supplementary Fig. 7b-d)**. For deeper selectivity profiling across the serine hydrolase family, we employed ABPP-ReDiMe in mouse brain proteomes as previously described^34^. ABD-110 potently inhibited AIG1 at both 1 and 10 mM with only a few off-targets detected, including ABHD6 and several highly homologous, mouse-specific carboxylesterases (CESs), which are common targets for carbamates^37^. The high potency of ABD-110 for AIG1 versus these off-targets demonstrates the high specificity of ABD-110 for AIG1; similar results were obtained in mouse kidney membranes (**Supplementary Fig. 7e-g**). AIG1 was not detected in these samples from chow-fed mouse WAT using untargeted MS-based approaches; as shown below, AIG1 was detected in mouse WAT in more directed target engagement studies.

**Fig. 8.**
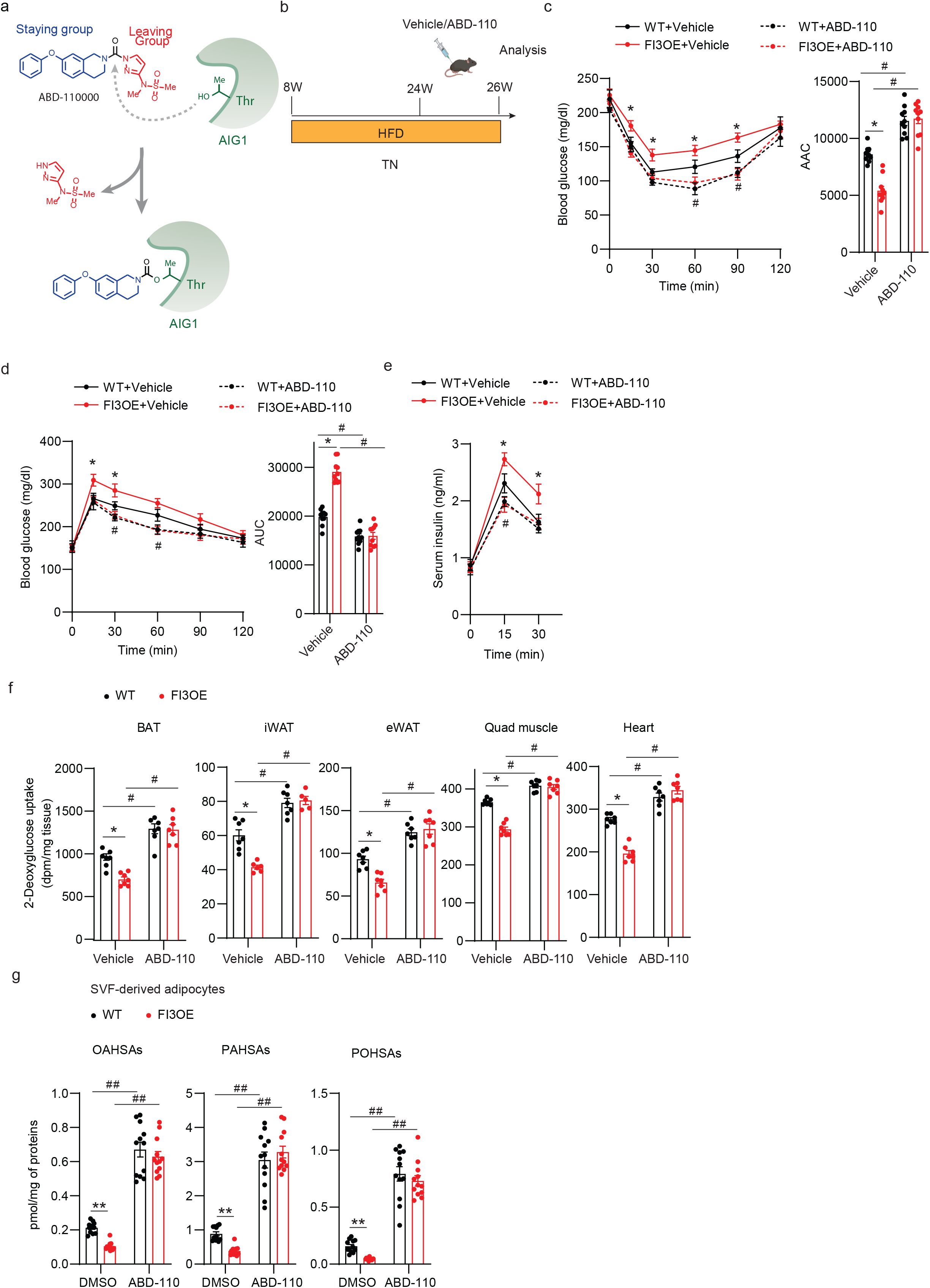
Pharmacologically inhibiting AIG1 attenuates HFD-induced insulin resistance in FI3OE mice. **a,** Proposed mechanism of covalent AIG1 inhibition by ABD-110. **b,** Schematic illustrating the strategy for the AIG1 inhibitor treatment of WT and FI3OE mice on HFD at thermoneutrality. Insulin tolerance test (**c**), glucose tolerance test (**d**), serum insulin levels (**e**), *in vivo* 2-deoxyglucose uptake in BAT, iWAT, eWAT, quadriceps muscle, and heart (**f**) of mice described in **b** (n=7-8). **g,** Quantification of FAHFA levels in WT and FI3OE SVF-derived adipocytes treated with or without ABD-110 (1μM, 4h) (n=12). Statistical significance was assessed by *two-way* ANOVA. Data in all panels are expressed as mean ± SEM. *P < 0.05 vs WT, **P < 0.01 vs WT, ^#^P < 0.05 vs. vehicle, ^##^P < 0.01 vs DMSO.

We next evaluated the potential of ABD-110 to be used *in vivo*. ABD-110 (2.5-25 mg/kg) was administered to C57/BL6J mice by oral gavage and, after 4 h, brains and kidneys were collected to evaluate AIG1 target engagement and selectivity by gel-based ABPP. Brain AIG1 inhibition could be observed at doses as low as 5 mg/kg with maximal inhibition being reached at 10 mg/kg. Notably, we did not observe significant off-target activity for other serine hydrolases or ADTRP with the exception of CES1c in the kidney which appeared to be maximally inhibited at 5 mg/kg (**Supplementary Fig. 7h, i)**. Finally, we assessed the effect of high fat feeding on the efficacy of the inhibitor. Chow or high fat-fed mice were dosed with vehicle or ABD-110 (25 mg/kg IP) once daily for 2 weeks. Four hours after the final dose, brain, kidney, eWAT and iWAT tissues were harvested for targeted MS-based ABPP analysis of AIG1 activity. In all tissues tested, the efficacy of AIG1 inhibition was excellent, with slightly improved results in high fat-fed mice (**Supplementary Fig. 7j, k)**. We speculate that ABD-110 may work better in the setting of obesity because endogenous FAHFAs are reduced^32^, providing less competition for binding to AIG1. Together these data support the utility of ABD-110 as a selective AIG1 tool for *in vivo* applications.

To assess the effect of ABD-110 on metabolic function, chow and high fat-fed WT mice were injected with vehicle or 25 mg/kg ABD-110 per day for 2 weeks. ABD-110 had no effect on insulin sensitivity and glucose tolerance on chow diet (**Supplementary Fig. 8a-c**). However, the impaired insulin and glucose tolerance of high fat-fed mice were ameliorated by ABD-110 treatment (**Supplementary Fig. 8d-f**). We next asked whether ABD-110 would rescue the metabolic dysfunction of FI3OE mice. High fat-fed WT and FI3OE mice were injected with vehicle or 25 mg/kg ABD-110 per day for 2 weeks, with no difference in body weight noted after ABD-110 injection (**Fig. 8b, Supplementary Fig. 9a, b**). The impaired insulin and glucose tolerance of high fat-fed FI3OE mice were ameliorated by AIG1 inhibition (**Fig. 8c, d**). Furthermore, the increased glucose-stimulated insulin secretion observed in FI3OE mice after glucose injection was reversed by ABD-110 treatment (**Fig. 8e**). Consistent with this, treatment with ABD-110 increased insulin-stimulated glucose uptake in all tissues tested in both WT and FI3OE mice and abolished the effects of adipocyte IRF3 overexpression (**Fig. 8f**). As before, FI3OE adipocytes exhibited reduced FAHFA levels. ABD-110 increased FAHFA levels in WT adipocytes and fully reversed the decrease in FI3OE adipocytes (**Fig. 8g, Supplementary Fig. 9c**).

## Discussion

Overnutrition and obesity promote a pro-inflammatory milieu in adipose tissue and other metabolically relevant tissues, and key extracellular mediators of inflammation, including cytokines like TNFα and IL-1β, and TLR ligands like LPS and cell-free DNA, are well established as endogenous drivers of insulin resistance^38–41^. Despite years of intensive study, there is still an incomplete understanding of how inflammatory signaling causes insulin resistance. Several proposed mechanisms involve direct cross-talk between inflammatory kinases like JNK and IKK and various insulin signaling components^42, 43^.

However, the time course over which insulin action is affected by inflammatory factors is much longer than would be expected if this were a primary mechanism^8, 24^. This suggests a potential role for the transcriptional effects of inflammation, a notion supported by the fact that alterations in chromatin state are known to affect insulin sensitivity, and also the proven utility of transcription factor-based insulin sensitizers^44, 45^. To date, however, few transcriptional pathways have been convincingly connected to insulin resistance. Among those that have been described, expression of *Slc2a4* (encoding the insulin-sensitive glucose transporter GLUT4) is reduced in adipocytes from insulin resistant humans, and targeted ablation of GLUT4 in adipocytes of mice resulted in insulin resistance in adipocytes, liver and muscle with reduced total body insulin sensitivity measured by euglycemic clamp studies^46^. In addition, expression of *Mlxipl* (encoding Carbohydrate Response Element Binding Protein, ChREBP) is reduced in adipose tissue of insulin resistant people^47^ and adipose specific knockout of ChREBP also results in insulin resistance in adipocytes, liver and muscle^48^. Finally, reduced *Pparg* expression has been noted in adipocytes from insulin resistant humans and rodents^46, 49^, but whether this causes insulin resistance is not known. We have provided evidence that the pro-inflammatory transcription factor IRF3 is a critical driver of insulin resistance in the context of obesity. IRF3 levels are increased in the adipocytes of people and mice with obesity, and mice lacking IRF3 in all tissues show improved insulin sensitivity before body weight divergence after high fat feeding^24^. We have also shown that IRF3 becomes activated in the hepatocytes and adipocytes of people and mice with obesity, where it mediates cellular and systemic insulin resistance^25, 26^. Interestingly, in hepatocytes, IRF3 drives insulin resistance by directly inducing the expression of *Ppp2r1b*, a component of the PP2A phosphatase complex^26^. The IRF3-mediated increase in PP2A activity diminishes AMPK and AKT phosphorylation, with subsequent impairment of glucose homeostasis. Here, we find that IRF3 is an inducer of *Aig1* in adipocytes, which reduces levels of insulin-sensitizing FAHFAs. Although the specific gene targets of IRF3 differ between these two cell types, both illustrate the key finding that pro-inflammatory factors ultimately affect insulin action through induction of genes that are outside the traditional inflammatory gene portfolio. Importantly, our data do not rule out potential roles for other downstream targets of IRF3, or for pro-inflammatory transcription factors besides IRF3 (AP-1 factors, for example). The novelty of the current work lies in (a) demonstrating that the adipocyte is a key cell type mediating the effect of IRF3 on insulin action, (b) showing definitively that the effect on insulin sensitivity is independent of weight changes, and most importantly, (c) identifying a specific molecular link between IRF3 and insulin resistance (i.e., reduction of FAHFAs by promoting expression of a FAHFA hydrolase).

FAHFAs have been shown to exert a myriad of beneficial metabolic actions, including improvement in insulin sensitivity and in glucose-stimulated insulin secretion. FAHFAs also exert anti-inflammatory effects, protecting islets from the effects of autoimmunity^50^ and also delaying the onset and reducing the severity of experimental colitis by modulating both innate and adaptive immune responses^51^. Our work illustrates that the converse is also true; inflammation leads to reduced FAHFA levels, at least in the context of overnutrition. Recently, ATGL has been shown to catalyze the formation of the FAHFA ester bond in mammals through its transacylase activity^52^. FAHFA degradation, on the other hand, is mediated by at least three hydrolases, AIG1 and ADTRP^34^, and carboxyl ester lipase^53^, the last of which is not expressed in adipocytes^54^. Insulin resistant mice and humans have reduced FAHFA levels in serum and subcutaneous adipose tissues^32^, although it has been unclear how this occurs. Here, we have shown that *Aig1* is a transcriptional target of IRF3 in adipocytes, and targeted lipidomic analysis from adipocytes identifies reductions in FAHFA levels as a consequence of IRF3-mediated inflammation. IRF3-mediated suppression of systemic glucose uptake was rescued by AIG1 inhibition, which is consistent with the previous finding that 5-PAHSA primes adipocytes for de novo lipogenesis^55^. and multiple FAHFA isomers directly augment insulin-stimulated glucose uptake in adipocytes^32, 56^ by augmenting GLUT4 translocation to the plasma membrane^32^. Interestingly, targeted ablation of AIG1 in all tissues failed to protect mice from HFD-induced insulin resistance and glucose intolerance^34^. The rationale for these conflicting results is unclear, and should be explored further.

## Methods

### Animal studies

All animal experiments were performed with approval from the Institutional Animal Care and Use Committees of the Harvard Center for Comparative Medicine and Beth Israel Deaconess Medical Center and Lundbeck La Jolla Research Center. Mice were maintained on a Chow diet (Harlan Teklad, 8664) under a 12h light/dark cycle at 22 °C for standard room temperature housing. Eight-week-old male mice were fed a 60% high-fat diet (D12492, Research Diets) in thermoneutrality for 16 weeks and body weight was measured weekly. All metabolic studies were performed in male mice. The IRF3 floxed mice (JAX:036260) and IRF3-2D mice (JAX:036261) were generated as previously described^25^. Strains purchased include C57BL/6 (JAX:000664) and *Adipoq-Cre* (JAX:028020).

### Local delivery of adeno-associated virus (AAV) in adipose tissues

Adeno-associated virus serotype 8 expressing mouse Aig1 and AAV8-GFP control were purchased from Applied Biological Materials (Canada). AAV injection was carried out as previously described^57^. Briefly, mice were anesthetized with an intraperitoneal injection of ketamine (100 mg/kg) and xylazine (10 mg/kg). A laparotomy was performed to expose eWAT. Each eWAT pad received four injections of 10 µL AAV solution using a Hamilton syringe to distribute the vector over the whole depot. The abdomen was rinsed with sterile saline solution and closed with a two-layer suture. Mice were monitored for changes in body weight and subjected to further analysis.

### Glucose tolerance tests (GTT) and Insulin tolerance tests (ITT)

For glucose tolerance tests (GTT), mice were fasted overnight and then injected intraperitoneally with glucose (1.5 g per kg body weight). For insulin tolerance tests (ITT), mice were fasted for 6 hrs and then injected intraperitoneally with insulin (Humulin, Eli Lilly; 1.5 U per kg body weight). Blood samples were collected from a tail nick at the indicated time points, and glucose levels were measured using test strips (OneTouch, USA).

### Serum Insulin Quantification

Blood samples from fed mice were centrifuged at 3,000 g for 10 min, and plasma was pipetted and stored at 80 °C. Serum insulin levels were measured using the ultra-sensitive mouse insulin ELISA kit according to the manufacturer’s instructions (Crystal Chem, USA).

### Mouse and human pre-adipocyte culture and differentiation

Inguinal subcutaneous WAT stromal-vascular fraction (SVF) cells were isolated from both male and female mice, cultured, and differentiated into mature adipocytes as previously described^58^. Briefly, inguinal white fat pads were obtained from male mice (4 to 6 weeks), and then dissected, washed three times with phosphate-buffered saline (PBS), minced into small pieces, and then digested for 45 mins in digestion buffer [1 mg/mL collagenase I (Sigma) and 1% bovine serum albumin (BSA) (Sigma-Aldrich) and 10mM CaCl_2_ (ThermoFisher) in PBS] at 37 °C. Digested tissues were passed through a sterile 100 µm cell strainer (BD Falcon) to remove undigested fragments and centrifuged at 600g for 5 min at 4°C to pellet the SVF cells. The pellet was resuspended in DMEM/F12+ GlutaMax (Invitrogen) medium containing 1% pen/strep and 10% FBS, then passed through a 40 µm cell strainer, centrifuged, resuspended as described above, and plated on collagen-coated culture dishes (Corning). For differentiation, SVF cells were grown to confluence and induced (day 0) in differentiation medium supplemented with 5 µ g/ml insulin (Sigma-Aldrich, I9278), 0.5mM 3-isobutyl-1-methylxanthine (IBMX, Sigma-Aldrich, I5879), 5 µM dexamethasone (Sigma-Aldrich, D4902), and 1 µM rosiglitazone (Sigma-Aldrich, R2408) for two days and subsequently cultured in maintenance medium supplemented with 5 µg/ml insulin and 1 µM rosiglitazone. On day 6, the SVF-derived adipocytes were transduced with 2 µl AAV-GFP or AAV-GFP-Cre for 12 h. Experiments were carried out on day 8 of differentiation.

Human SGBS preadipocytes were cultured and differentiated as previously described^29^ with slight modifications. SGBS preadipocytes were seeded in a 12-well plate at 8×10^4^ per well and cells were cultured in a normal culture medium (DMEM/F12+ GlutaMax with 1% pen/strep and 10% FBS) until they reached confluency. Cells were incubated in normal culture medium (DMEM/F12+ GlutaMax with 1% pen/strep and 10% FBS) with 4 ng/ml human FGF (Perprotech, 100-18B) for 2 days, then incubated (day 0) in differentiation medium [1% pen/strep, 33 nM Biotin (Sigma-Aldrich, B-4639), 17 nM Panthotenate (Sigma-Aldrich, P-5155), 10 µg/ml Transferrin (Sigma-Aldrich, T-2252), 20nM insulin, 100nM cortisol (Sigma-Aldrich, H-0888), 0.2nM T3 (Sigma-Aldrich, T-6397), 25nM dexamethasone, 250µM IBMX and 2 µM rosiglitazone in DMEM/F12+ GlutaMax] for 4 days. After the fourth day, the cells were incubated in the maintenance medium (1% pen/strep, 33 nM, 17 nM Panthotenate, 10 µg/ml Transferrin, 20nM insulin, 100nM cortisol, and 0.2nM T3 in DMEM/F12+ GlutaMax); maintenance medium was changed every fourth day. Experiments were carried out on days 10-day and 12.

### RNA-Seq

For RNA-seq analysis, sequencing reads were demultiplexed and trimmed for adapters using bcl2fastq (v2.20.0). Secondary adapter trimming, NextSeq/Poly(G) tail trimming, and read filtering was performed using fastp (v0.20.1)^59^; low quality reads and reads shorter than 24nt after trimming were removed from the read pool. Salmon (v1.4.0)^60^ was used to simultaneously map and quantify reads to transcripts in the GENCODE M24 genome annotation of GRCm38/mm10 mouse assembly. Salmon was run using full selective alignment, with sequence-specific and fragment GC-bias correction turned on (--seqBias and --gcBias options, respectively). Transcript abundances were collated and summarized to gene abundances using the tximport package for R^61^. Normalization and differential expression analysis were performed using edgeR. For differential gene expression analysis, genes were considered significant if they passed a fold change (FC) cutoff of log2FC≥1 and a false discovery rate (FDR) cutoff of FDR≤0.05. This paper’s accession number for the RNA-seq datasets is GEO: GSE213048.

### RNA Isolation/Quantitative RT-PCR

Total RNA was extracted from cells or tissues using a Direct-zol RNA MiniPrep kit (ZYMO Research, USA) or Trizol reagent (Invitrogen, USA). Reverse transcription was performed with 1µg of total RNA using a High-Capacity cDNA Reverse Transcription Kit (Thermo Fisher, USA). qRT-PCR was performed on an ABI PRISM 7500 (Applied Biosystems, USA). Melting curve analysis was carried out to confirm the RT-PCR products. Statistical analysis was performed using the ΔΔCT method with TBP primers as control (primer sequences in **Extended Data Table 2**).

### Immunoblotting

For immunoblotting analyses, tissues and cells were lysed in RIPA buffer containing protease and phosphatase inhibitors (Thermo Fisher, USA). Protein levels were quantified using a BCA protein assay kit (Thermo Fisher, USA) and lysates containing an equal amount of protein were subjected to SDS-PAGE and transferred to polyvinylidene fluoride (PVDF) membranes, followed by incubations with primary and secondary antibodies. Primary antibodies against phospho AKT(Ser473) (CST, 9271), AKT (AKT, 4685), HSP90 (CST, 4877), phospho IRF3(Ser396) (CST, 29047), IRF3 (CST, 11904) and AIG1 (Proteintech, 14468-1-AP), were used.

### Glucose uptake assays

Glucose uptake assays were performed as described earlier^24^. Briefly, SVF-derived adipocytes (day 6) and human SGBS cells (day 14) were incubated in serum-free DMEM/F12 for 4–6 hours. Cells were then washed 3 times with KRH buffer (12 mM Hepes, pH 7.4, 121 mM NaCl, 5 mM KCl, 0.33 mM CaCl_2_, and 1.2 mM MgSO_4_) and incubated for 20 minutes in KRH buffer in the absence or presence of 100 nM insulin. Cells were treated with 2-deoxy-d-[2,6-^3^H]-glucose (0.33 μCi/ml) for another 10 minutes. Glucose uptake was stopped quickly by 3 rapid washes with KRH buffer containing 200 mM glucose and 10 μM cytochalasin B on ice. Cells were solubilized in 0.1% SDS for 30 minutes, and radioactivity was measured by liquid scintillation counting. In some experiments, a combination of 9-POHSA (20µM), 9-PAHSA (20µM) and 9-OAHSA (20µM) (Cayman Chemical), or Vehicle (DMSO, 0.01%) were incubated with SVF-derived adipocytes for 24 hours, then insulin-stimulated glucose uptake studies were initiated. Total protein was determined by the BCA method (Pierce), and results were normalized to protein amount.

### Insulin signaling *in vivo*

Mice were fasted overnight for insulin signaling studies, and insulin (10 U/kg body weight) was administered i.p. After 10 minutes various tissues were harvested and stored at –80°C until use. Tissue samples were homogenized in cell signaling lysis buffer containing protease inhibitors (Roche) and phosphatase inhibitors (Sigma-Aldrich) and subjected to Western blotting.

### Construct design and site-directed mutagenesis

To construct the mouse AIG1 overexpression lentivirus, the open reading frames of mouse *Aig1* was cloned into pCDH-CMV-MCS-EF1-Puro cDNA Cloning and Expression Vector. To construct the shAIG1 lentivirus, the short hairpins (5’-ACCTTCTCCGTGGGCTATATA-3’ and 5’-CTATGACAGAGAGATGATATA-3’) were cloned into the pLKO.1-TRC cloning vector, respectively. To construct shIRF3 lentivirus, the short hairpin (5’-CGAAGTTATTTGATGGCCTGA-3’) was cloned into the pLKO.1-TRC cloning vector. To generate dox-inducible IRF3 overexpression lentivirus, mouse IRF3 cDNA was cloned into pCW57-MCS1-P2A-MCS2 (Hygro). To construct dox-inducible shIRF3 lentivirus, the short hairpin (5’-CGAAGTTATTTGATGGCCTGA-3’) was cloned into the Tet-pLKO-puro cloning vector. Mutagenesis primers used to construct AIG1(T43A) were as follows: forward, 5’-GAAGTTCCTGGCCTTCATTGATC-3’; reverse, 5’-GATCAATGAAGGCCAGGAACTTC-3’; mutagenesis primers for AIG1(H134A): forward, 5’-CCATGGAATGGCCACAACGGTTT-3’; reverse, 5’-AAACCGTTGTGGCCATTCCATGG-3’.

### Lentivirus packaging

psPAX2 and pMD2.G were used for the lentiviral packaging. High-titer lentivirus was packaged in 293T cells using lipofectamine 2000 transfection agent. Viral supernatants were collected 48h post-transfection and used for pre-adipocyte infection.

### Tissue glucose uptake *in vivo*

In vivo tissue glucose uptake studies were carried out with mice maintained on HFD in thermoneutrality. After a 4-hour fast, mice were injected with insulin (Humulin, 0.75 U/kg, ip) and a trace amount of 2-deoxy-D-[1-^14^C] glucose (10 µCi; PerkinElmer). Forty minutes later, the mice were euthanized, and tissues were collected, weighed, and homogenized. Radioactivity was measured and counted as described^62^.

### Lipid extraction for FAHFA analysis

For the measurement of FAHFAs from SVF-derived cells, total lipids were extracted using the modified Bligh–Dyer method^63^. In brief, 1.5 ml of 2:1 chloroform/methanol with the internal standards [^13^C_16_] 9-PAHSA, [^13^C_18_] 12-OAHSA, and [^13^C_16_] 5-PAHSA was added to 500 µl of cell suspension in PBS. Samples were vortexed and centrifuged at 2,000g for 7 min. The bottom organic phase was transferred into a new vial and dried under a N_2_ stream.

eWAT samples (50 mg) were dounce homogenized in the same mixture of solvents described above and then total lipids were extracted with the same method used for SVF-derived cells. FAHFAs were enriched using a solid-phase extraction (SPE) column (Hypersep silica 500 mg) as described previously^64^. Briefly, columns were equilibrated with 15 ml hexane, and the samples were then resuspended in 200 µl chloroform and loaded on the SPE column. The neutral lipid fraction containing TGs was eluted with 16Lml of 95:5% hexane/ethyl acetate, and the polar lipid fraction containing non-esterified FAHFAs was eluted using 15 ml ethyl acetate. The FAHFA fractions were dried under a stream of N_2_ and stored at −40 °C until LC-MS/MS analysis.

### FAHFA analysis using LC–MS/MS

FAHFA isomers were quantified using an Agilent 6470 Triple Quad LC–MS/MS instrument through multiple reaction monitoring (MRM) in the negative ionization mode as previously described^52^. In brief, cell culture lipid extracts were reconstituted in 50 µl methanol, and tissue extracts were reconstituted in 100 µl methanol. 7 µl of each sample were injected onto a UPLC BEH C18 Column (Waters Acquity, 186002352). FAHFA regioisomers were resolved using 93:7 methanol/water with 5 mM ammonium acetate and 0.01% ammonium hydroxide solvent through an isocratic gradient at 0.15LmlLmin^−1^ flow rate for 60Lmin.

### ABD-110 In vivo Dose-Response

Female C57Bl/6J mice (Jackson) aged 10 weeks at the time of dosing were administered ABD-110 by oral gavage (10 ml/kg volume). ABD-110 was prepared fresh on the day of dosing in water containing 5% solutol and 20% HpbCD. Maximal dispersal of the compound was achieved by bath and probe sonication until a uniform white suspension was formed. Animals were administered single oral doses of ABD-110 (2.5–25 mg/kg). Four hours after compound administration, animals were anesthetized with isoflurane and decapitated. Brains and kidneys were removed and rinsed in PBS before freezing in liquid nitrogen. Tissues were stored at -80 °C until analysis by gel-based ABPP (see below).

## Proteome Preparation for ABPP

### Mouse tissue (brain, kidney and adipose) homogenates

Frozen mouse tissues (one brain hemisphere, one whole kidney or ∼100 mg of WAT) were homogenized by adding the tissue into a 2 mL Safe-lock microcentrifuge tube along with 0.5 -1.0 mL of PBS and a 5 mm steel bead. The tissue was then homogenized using the Qiagen TissueLyzer at 30 Hz for 1 min. After homogenization, samples were put through a clarifying spin (1,000 *g*, 10 min, 4 °C) and supernatants were collected.

Brain and kidney homogenates were fractionated further by ultra-centrifugation (100,000 *g*, 45 min, 4°C) and membrane pellets were resuspended in PBS. Protein concentrations were determined using Bio-Rad DC protein assay and diluted to desired protein concentration (see below) for subsequent ABPP analysis.

## Determination of ABD-110 Potency and Selectivity by ABPP

### Gel-based ABPP

Inhibitor potency (IC_50_ values and target engagement) against AIG1 and other serine/threonine hydrolases was determined by competitive gel-based ABPP in mouse brain and kidney membrane homogenates using FP-Rh competition^65^. Mouse brain and kidney proteomes were prepared as described above. For determination of mADTRP activity, full length mADTRP (Dharmacon) was recombinantly expressed in HEK293T cells and the corresponding cell lysates were doped into mouse brain membrane proteomes (1.0 mg/mL) at a ratio to enable detection of both AIG1 and ADTRP.

*In vitro* potency for AIG1, ADTRP and other serine hydrolase enzymes was determined by treating brain proteomes (50 µg, 1.0 mg/mL with or without mADTRP-containing cell lysates) with ABD-110 or DMSO for 30 min at 37°C and subsequently treated with FP-Rh (1.0 µM) for an additional 30 min at room temperature.

For gel-based *in vivo* target engagement, brain and kidney proteomes (50 µg) from vehicle or ABD-110-treated mice were treated with FP-Rh (1.0 µM) for 30 min at room temperature.

After incubation with FP-Rh, reactions were quenched with 4X SDS-PAGE loading buffer and FP-Rh-labeled enzymes were resolved by SDS-PAGE (10% acrylamide). In-gel fluorescence was visualized using a Bio-Rad ChemiDoc^TM^ XRS imager. Fluorescence is shown in gray scale. Quantification of enzyme activities was performed by densitometric analysis using ImageJ software (NIH). Integrated peak intensities were generated for the band corresponding to AIG1^34^. IC_50_ values were calculated through curve fitting semi-log-transformed data (*x*-axis) by non-linear regression with a four-parameter, sigmoidal dose response function (variable slope) in Prism software (GraphPad).

### MS-based ABPP

Brain, kidney and adipose tissue proteomes (2.5 mg/mL in 0.2 mL of PBS) were treated with inhibitor at the specified concentrations (0.001-10 μM) or DMSO for 30 min at 37 °C and subsequently labeled with FP-biotin (10 μM) for 1 h at room temperature. Proteome samples were immediately precipitated and processed through the MS-based ABPP sample preparation protocol (below).

### MS-based ABPP Sample Preparation

Tissue proteomes (2.5 - 5 mg/mL normalized by tissue source) in 200 mL of PBS were labeled with FP-biotin (50 μM) for 1 h at 37 °C. After labeling, the proteomes were denatured and precipitated using 9:1 acetone/MeOH, resuspended in 0.2 mL of 8 M urea in PBS and 1% SDS, reduced using DL-dithiothreitol (DTT, 10 mM) for 20 min at 55 °C, and then alkylated using iodoacetamide (50 mM) for 30 min at room temperature in the dark. The biotinylated proteins were enriched with PBS-washed streptavidin-agarose beads (50 μL; Thermo Scientific) by rotating at room temperature for 1.5 h in PBS with 0.2% SDS (1.3 mL). The beads were then washed sequentially with 0.5 mL 0.2% SDS in PBS (10×), 1.1 mL PBS (10×) and 1.1 mL DI H_2_O (10×). On-bead digestion was performed using sequencing-grade trypsin (2 μg; Promega) in 2 M urea in PBS for 12–14 h at room temperature (100 μL). The beads were removed using filtration and washed with DPBS (100 µL).

For *in vitro* MS-based ABPP selectivity profiles in mouse brain and kidney, peptide digests derived from vehicle (DMSO) and ABD-110-treated proteomes were dimethylated with light (CH_2_O) or heavy (^13^CD_2_O) formaldehyde and NaBH_3_CN as previously described^66^. After 1 h, the methylation reaction was quenched with NH_4_OH (40 μL, 1%) and, subsequently, formic acid (20 μL, 100%) before the heavy and light samples were combined. The mixed samples were then desalted as described below. For all other samples, the digests were acidified by addition of formic acid (10 µL of 100% formic acid) and desalted using SOLAµ^TM^ SPE Plates (HRP 2 mg / 1 mL). Samples were dried by centrifugal evaporation. *In vitro* ABD-110-treated WAT samples were labeled with TMTpro reagents (see below) whereas *in vivo* ABD-110-treated samples were stored at -80 °C until targeted MS analysis.

### TMTpro Labeling Protocol

TMTpro labeling protocols were adapted from previous work^67^. Dried samples were reconstituted in 20 µL EPPS buffer, pH 8.5 followed by addition of 5 µL 40 mM TMTpro reagents in acetonitrile for 1 hr. Reactions were quenched with 5 µL 5% hydroxylamine, mixed, desalted, dried as in prior section, and stored at -80 °C until analysis.

## MS Data Acquisition and Analysis

### Parallel reaction monitoring (PRM) for in vivo target engagement

Dry peptide samples were reconstituted in water containing 0.1% formic acid (20 µL) and 10 µL were injected onto an EASY-Spray column (15 cm x 75 µm ID, PepMap C18, 3 µm particles, 100 Å pore size, Thermo Fisher Scientific) using a Vanquish Neo UHPLC (Thermo Fisher Scientific). Peptides were separated over a 15 min gradient of 0 to 40% acetonitrile (0.1% formic acid) and analyzed on an Orbitrap Fusion Lumos (Thermo Fisher Scientific) operated using a parallel reaction monitoring (PRM) method for two AIG1 peptides and additional control peptides to assess sample integrity: FASN, PCCA, PC. Selected ions were isolated and fragmented by high energy collision dissociation (HCD) at 30% CE and fragments were detected in the Orbitrap at 15,000 resolution. Further details for the targeted peptides can be found below.

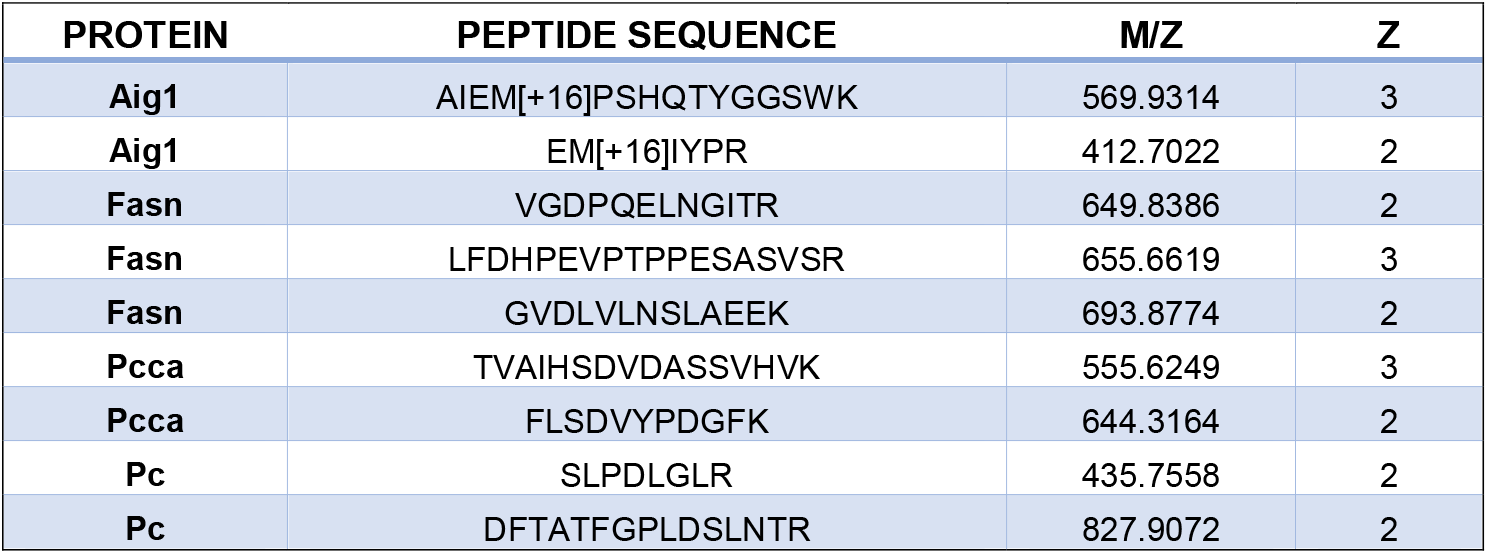

### MS Acquisition for TMTpro Labeled Peptides

MS was performed using a Thermo Scientific Vanquish Neo and Orbitrap Eclipse system. Peptides were eluted over a 240 min nLC 5-25-45% acetonitrile gradient. For all samples, data were collected in data-dependent acquisition mode over a range from 375–1,500 m/z. Each full scan was followed by fragmentation events for 1.5 sec including real-time search of mouse peptides with TMTpro modifications and carbamidomethylation of cysteine and oxidation of methionine for subsequent synchronous precursor selection. Dynamic exclusion was enabled (repeat count of 1, exclusion duration of 60 s) for all experiments.

### Reductive Dimethylation Data Analysis

Data analysis was performed using a custom in-house pipeline. Raw files were converted, searched and processed with the same software as described for the TMTpro datasets, with the following modifications. Comet searches were carried out with static modifications for carbamidomethylation at cysteine (+57.021463 Da), and the light dimethyl label at lysine and the N-terminus (+28.031300 Da), and differential modifications for the heavy dimethyl label at lysine and the N-terminus (+6.031817 Da) and for methionine oxidation (+15.994914 Da). Peptide ratio quantification was then performed using CIMAGE^68^. Competition ratios for each were then calculated from the median of all peptide ratios, and then normalized so that the median protein ratio for each sample was equal to one, capped to a maximum absolute value of 32, and converted to percent competition values. Percent competition values for serine hydrolases were reported only if the protein had at least 3 quantified peptides, and at least 1 unique quantified peptide, with an exception being made for AIG1, which was included regardless of its numbers of quantified peptides.

### TMTpro Data Analysis

Data analysis was performed using a custom in-house pipeline. Raw files were converted to the indexed mzML format using ThermoRawFileParser v1.4.2^69^, and then searched with Comet v2019.01.5^70^ against a human FASTA database obtained from UniProt^71^ (reference proteome UP000005640_9606, downloaded on 2020-03-21), and concatenated with a list of non-human contaminant proteins obtained from MaxQuant^72^, and reverse decoys. For the Comet search, TMTPro modifications on lysine and the N-terminus were specified as static modifications, in addition to a static modification for carbamidomethylation at cysteine [+57.021463], and a differential modification for oxidation at methionine [+15.994914]. Further processing was performed using the OpenMS platform v3.0.0^73^ (commit short hash 26292ad). Files were first converted from .pep.xml to .idXML using IDFileConverter, target/decoy information was added using PeptideIndexer, features were extracted with PSMFeatureExtractor, and Percolator v3.05^74^ was run through PercolatorAdapter. Peptide spectrum matches (PSMs) were then filtered with a q-value threshold of 0.01. Further processing was then carried out using custom Python code. For each filtered PSM, MS3 scans were extracted from the original mzML file, and reporter ion intensities were extracted from the most intense peak within a 0.002 Th window of each reporter ion expected mass. Intensity values were then normalized per channel to the mean per-channel summed intensity. PSMs were excluded from downstream calculations if their control sum intensity was less than 5000 x N where N is the number of channels corresponding to the control condition. Intensity values for each PSM were then summed per-channel for each protein, and percent activity values were calculated by dividing each value by the mean intensity across the summed control channels. The mean was then taken across channels associated with the same condition and was reported as a percent activity value for each protein.

## Chemistry

### Synthesis of ABD-11000

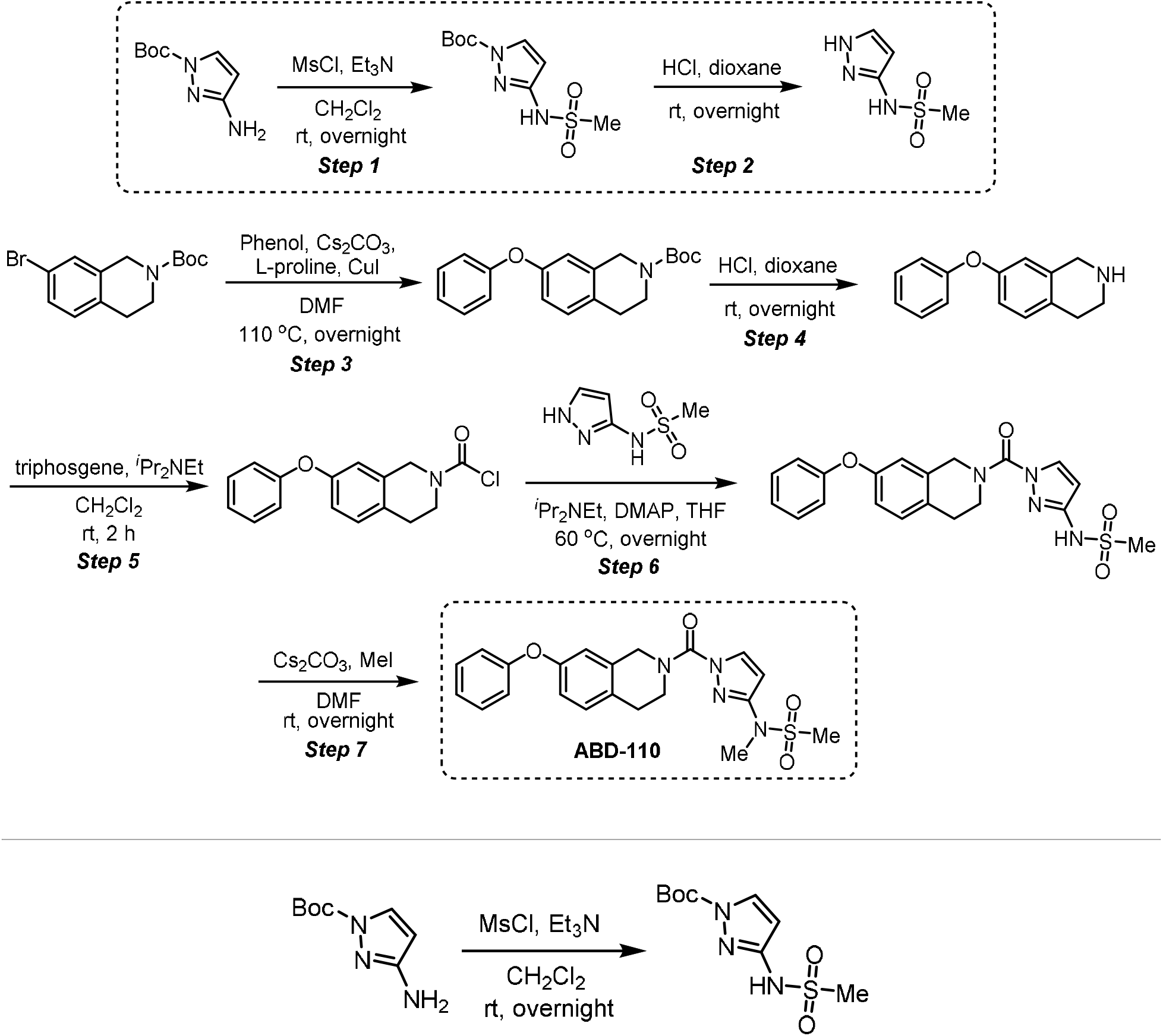

**Step 1 - *tert*-butyl 3-(methylsulfonamido)-1H-pyrazole-1-carboxylate**: A 100-mL round-bottom flask was charged with tert-butyl 3-amino-1H-pyrazole-1-carboxylate (300 mg, 1.64 mmol, 1.00 equiv), methanesulfonyl chloride (207 mg, 1.81 mmol, 1.10 equiv), dichloromethane (10 mL) and triethylamine (497 mg, 4.91 mmol, 3.00 equiv). The resulting solution was stirred overnight at room temperature and quenched by water (40 mL). The resulting solution was extracted with dichloromethane (3 x 80 mL) and the organic layers were combined, washed with water (3 x 20 mL), dried over anhydrous sodium sulfate, filtered and concentrated under reduced pressure to provide 400 mg (crude) of *tert*-butyl 3-(methylsulfonamido)-1H-pyrazole-1-carboxylate which was carried to the next step without further purification. LCMS (ESI, *m/z*): 262 [M+H]^+^.

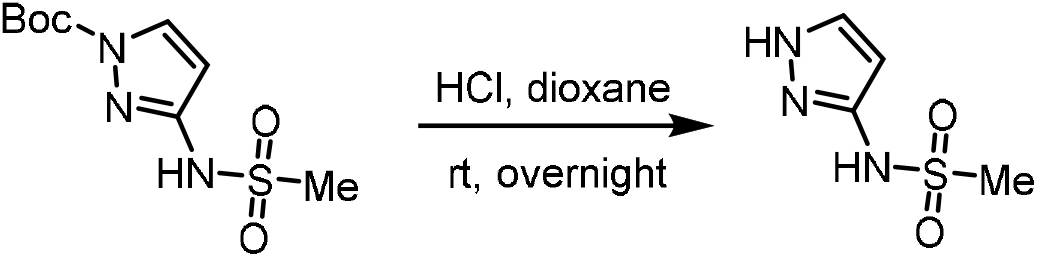

**Step 2 - N-(1H-pyrazol-3-yl)methanesulfonamide:** A 100-mL round-bottom flask was charged with *tert*-butyl 3-(methylsulfonamido)-1H-pyrazole-1-carboxylate (400 mg, 1.53 mmol, 1.00 equiv), 1,4-dioxane (10 mL) and concentrated hydrogen chloride (5 mL). The resulting solution was stirred overnight at room temperature and concentrated under reduced pressure to provide 450 mg (crude) of N-(1H-

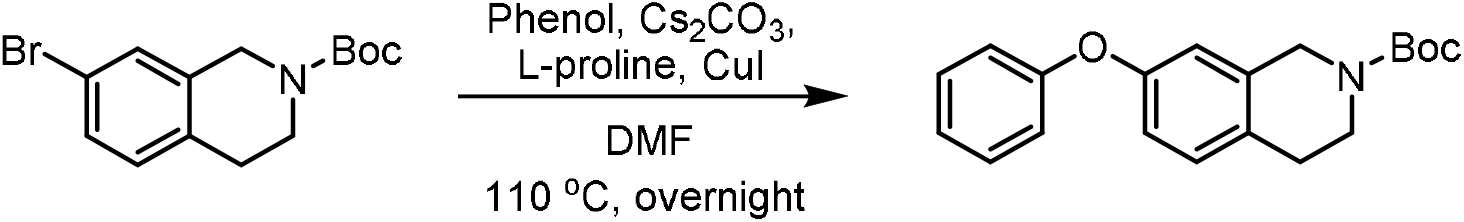

**Step 3 - *tert*-butyl 7-phenoxy-3,4-dihydroisoquinoline-2(1H)-carboxylate:** A 100-mL round-bottom flask was charged with *tert*-butyl 7-bromo-1,2,3,4-tetrahydroisoquinoline-2-carboxylate (2.00 g, 6.41 mmol, 1.00 equiv), *N,N*-dimethylformamide (50 mL), phenol (1.20 g, 12.7 mmol, 2.00 equiv), dicesium carbonate (5.20 g, 16.0 mmol, 2.50 equiv), L-proline (298 mg, 2.57 mmol, 0.40 equiv) and cuprous iodide (244 mg, 1.28 mmol, 0.20 equiv) under nitrogen. The resulting solution was stirred overnight at 110 °C and quenched by water (40 mL). The resulting solution was extracted with dichloromethane (3 x 80 mL) and the organic layers were combined, washed with water (3 x 20 mL), dried over anhydrous sodium sulfate, filtered and concentrated under reduced pressure. The residue was chromatographed on a silica gel column with ethyl acetate/petroleum ether (1/10) to provide 1.25 g (60% yield) of *tert*-butyl 7-phenoxy-1,2,3,4-tetrahydroisoquinoline-2-carboxylate as light yellow oil.

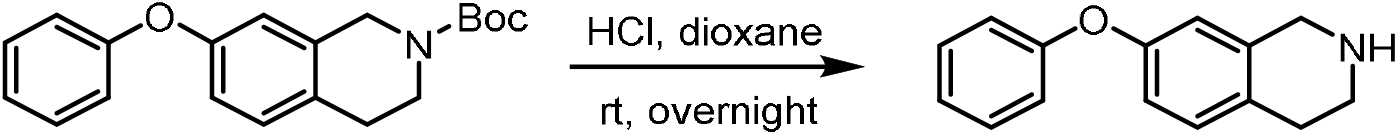

**Step 4 - 7-phenoxy-1,2,3,4-tetrahydroisoquinoline:** A 50-mL round-bottom flask was charged with *tert*-butyl 7-phenoxy-1,2,3,4-tetrahydroisoquinoline-2-carboxylate (1.25 g, 3.84 mmol, 1.00 equiv), 1,4-dioxane (10 mL), concentrated hydrogen chloride (5 mL). The resulting solution was stirred overnight at room temperature and concentrated under reduced pressure. The resulting solution was diluted with water (40 mL). The pH value of the solution was adjusted to 11-12 with sodium hydroxide (4 M). The resulting solution was extracted with dichloromethane (3 x 80 mL) and the organic layers were combined, washed with water (3 x 20 mL), dried over anhydrous sodium sulfate, filtered and concentrated under reduced pressure to provide 1.10 g (crude) of 7-phenoxy-1,2,3,4-tetrahydroisoquinoline as light yellow oil. LCMS (ESI, *m/z*): 226 [M+H]^+^.

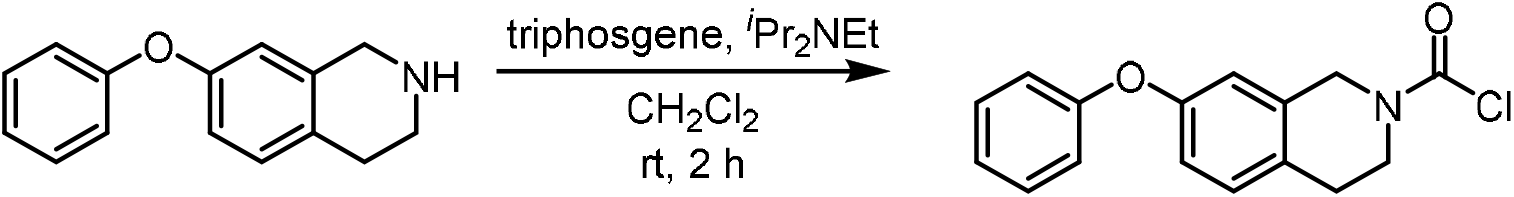

**Step 5 - 7-phenoxy-3,4-dihydroisoquinoline-2(1H)-carbonyl chloride:** A 50-mL round-bottom flask was charged with triphosgene (158 mg, 0.530 mmol, 0.50 equiv), dichloromethane (5 mL), 7-phenoxy-1,2,3,4-tetrahydroisoquinoline (240 mg, 1.07 mmol, 1.00 equiv). *N,N*-diisopropylethylamine (413 mg, 3.20 mmol, 3.00 equiv) was added dropwise at 0 °C. The resulting solution was stirred for 2 h at room temperature and quenched by water (40 mL). The resulting solution was extracted with dichloromethane (3 x 80 mL) and the organic layers were combined, washed with water (3 x 20 mL), dried over anhydrous sodium sulfate, filtered and concentrated under reduced pressure to provide 300 mg (crude) of 7-phenoxy-1,2,3,4-tetrahydroisoquinoline-2-carbonyl chloride as light yellow oil. This was used in the next step without further purification.

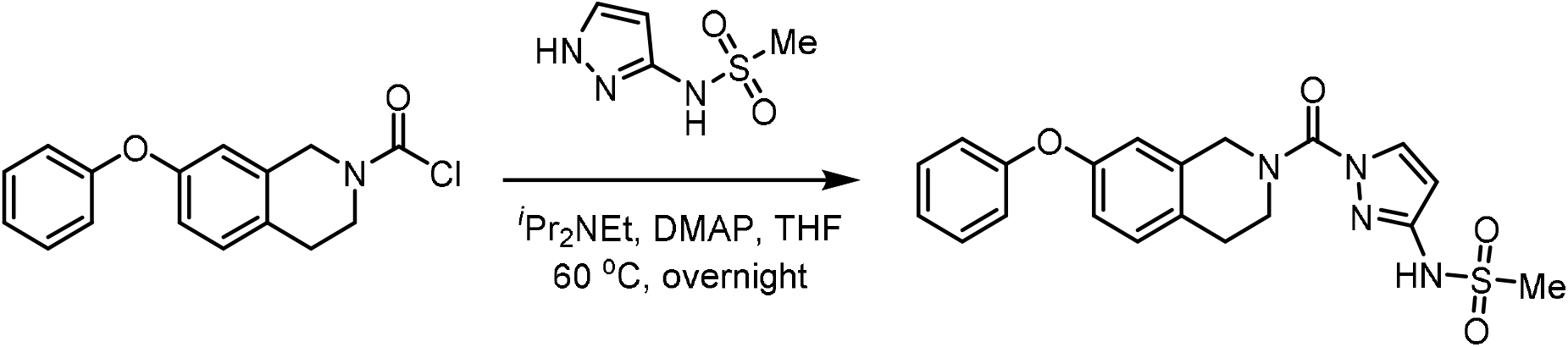

**Step 6 - N-(1-(7-phenoxy-1,2,3,4-tetrahydroisoquinoline-2-carbonyl)-1H-pyrazol-3-yl)methanesulfonamide:** A 50-mL round-bottom flask was charged with 7-phenoxy-1,2,3,4-tetrahydroisoquinoline-2-carbonyl chloride (150 mg, 0.520 mmol, 1.00 equiv), tetrahydrofuran (5 mL), *N*-(1H-pyrazol-3-yl)methanesulfonamide (84.0 mg, 0.520 mmol, 1.00 equiv), *N,N*-diisopropylethylamine (202 mg, 1.56 mmol, 3.00 equiv) and 4-dimethylaminopyridine (13.0 mg, 0.110 mmol, 0.20 equiv). The resulting solution was stirred overnight at 60 °C and quenched by water (40 mL). The resulting solution was extracted with dichloromethane (3 x 80 mL) and the organic layers were combined, washed with water (3 x 20 mL), dried over anhydrous sodium sulfate, filtered and concentrated under reduced pressure. The residue was chromatographed on a silica gel column with ethyl acetate/petroleum ether (1/1) to provide 170 mg (79% yield) of *N*-[1-[(7-phenoxy-1,2,3,4-tetrahydroisoquinolin-2-yl)carbonyl]-1H-pyrazol-3-yl]methanesulfonamide as light yellow oil. LCMS (ESI, *m/z*): 413 [M+H]^+^.

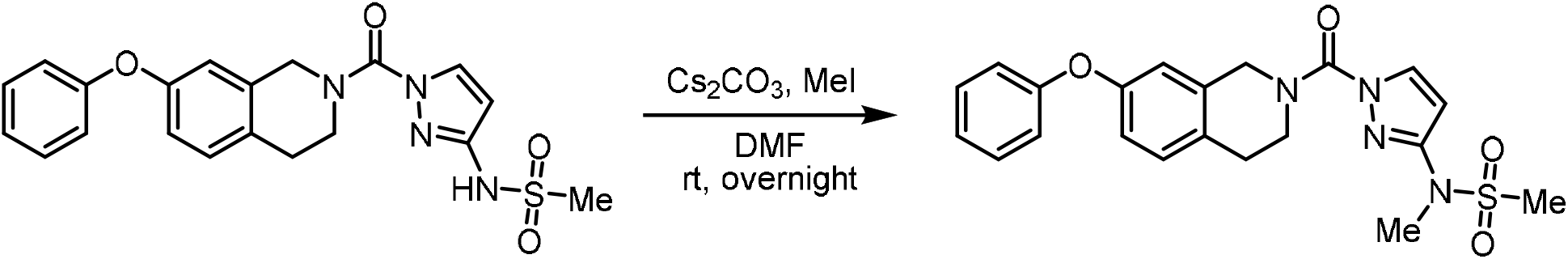

**Step 7 - N-methyl-N-(1-(7-phenoxy-1,2,3,4-tetrahydroisoquinoline-2-carbonyl)-1H-pyrazol-3-yl)methanesulfonamide (ABD-110):** A 50-mL round-bottom flask was charged with *N*-[1-[(7-phenoxy-1,2,3,4-tetrahydroisoquinolin-2-yl)carbonyl]-1H-pyrazol-3-yl]methanesulfonamide (170 mg, 0.410 mmol, 1.00 equiv), *N,N*-dimethylformamide (4 mL), dicesium carbonate (269 mg, 0.830 mmol, 2.00 equiv) and iodomethane (88.0 mg, 0.620 mmol, 1.50 equiv). The resulting solution was stirred overnight at room temperature and quenched by water (40 mL). The resulting solution was extracted with dichloromethane (3 x 80 mL) and the organic layers were combined, washed with water (3 x 20 mL), dried over anhydrous sodium sulfate, filtered and concentrated under reduced pressure. The crude product was purified by preparative HPLC using the following gradient conditions: 20% CH_3_CN/80% Phase A increasing to 80% CH_3_CN over 10 min, then to 100% CH_3_CN over 0.1 min, holding at 100% CH_3_CN for 1.9 min, then reducing to 20% CH_3_CN over 0.1 min, and holding at 20% for 1.9 min, on a Waters 2767-5 Chromatograph. Column: Xbridge Prep C_18_, 19*150mm 5um; Mobile phase: Phase A: aqueous NH_4_HCO_3_ (0.05%); Phase B: CH_3_CN; Detector, UV220 & 254nm. Purification resulted in 103 mg (58% yield) of *N*-methyl-N-[1-[(7-phenoxy-1,2,3,4-tetrahydroisoquinolin-2-yl)carbonyl]-1H-pyrazol-3-yl]methanesulfonamide as a light yellow semi-solid. ^1^H NMR (400 MHz, CDCl_3_) δ 8.07 (d, *J* = 2.8 Hz, 1H), 7.31 - 7.36 (m, 2H), 7.09 - 7.16 (m, 2H), 6.98 - 7.00 (m, 2H), 6.87 - 6.89 (m, 1H), 6.77 (s, 1H), 6.52 (d, *J* = 2.8 Hz, 1H), 4.89 (s, 2H), 4.03 (s, 2H), 3.39 (s, 3H), 3.02 (t, *J* = 5.8 Hz, 2H), 2.94 (s, 3H). LCMS (ESI, *m/z*): 427 [M+H]^+^.

### Statistics and reproducibility

All data are presented as mean ± s.e.m. Statistical analysis was performed in Prism 8 (GraphPad Software). Comparisons between two groups were performed using an unpaired two-tailed Student’s *t-test*. A *one-way* ANOVA followed by Dunnett’s or Bonferroni’s multiple comparisons test was used to compare more than two groups containing one factor. A *two-way* ANOVA followed by Dunnett’s multiple comparisons test was used to compare more than two groups containing two factors. P < 0.05 was considered to be statistically significant.

### Reporting summary

Further information on research design is available in the Nature Research Reporting Summary linked to this article.

### Data availability

There are no restrictions as to the availability of materials reported in the manuscript. The data that support the findings of this study are available from the corresponding author upon reasonable request. The RNA-seq data generated in this study have been deposited in the NCBI Gene Expression Omnibus and are accessible through accession number GSE21304. Source data are provided with this paper.

## Acknowledgements

This work was supported by NIH R01 DK102170 to E.D.R. and by support from the Dasman Diabetes Institute/Kuwaiti Ministry of Health to E.D.R. and R.A, NIH grants R01 DK43051 to B.B.K. and R01 DK106210 to B.B.K. and A.Sag., and a grant from the JPB Foundation to B.B.K; NIH K01 128075 to A. San. We thank the NIH BNORC supported Functional Genomics and Bioinformatics Core at BIDMC. This work was supported by the Mass Spectrometry Core of the Salk Institute with funding from NIH-NCI CCSG: P30 014195 and the Helmsley Center for Genomic Medicine. The MS data for untargeted lipidomics was gathered on a Thermo Fisher Q Exactive Hybrid Quadrupole Orbitrap mass spectrometer funded by NIH grant (1S10OD021815-01). We also thank Kerry Wellenstein for technical help.

## Author contributions

Conceptualization, S.Y., and E.D.R.; Investigation, S.Y., A.San, N.N., R.M.S., B.R.F., R.B.H., C.L.H., D.M.H., M.J.N., A.M.P; Data analysis, C.L.J., S.Y.; Writing, S.Y., and E.D.R.; Funding Acquisition, B.B.K., R.A., E.D.R.; Supervision and data interpretation, A.Sag, B.B.K., R.A., and E.D.R.

---

**Supplemental Fig. 1.**
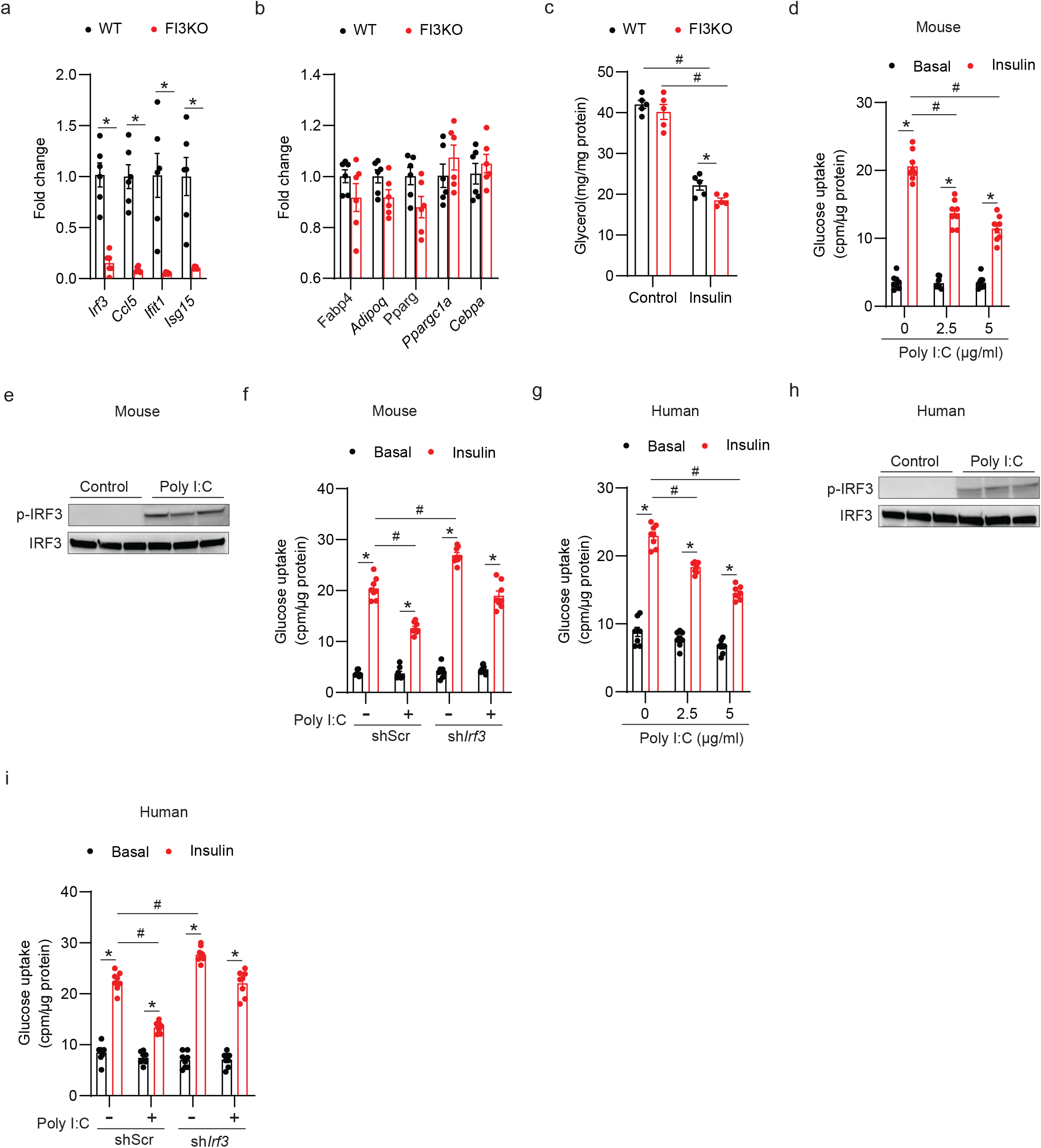
IRF3 is necessary for TLR3-mediated insulin resistance in mouse and human adipocytes. **a,** mRNA analysis of *Irf3*, *Ccl5*, *Ifit1*, and *Isg15* in WT and FI3KO SVF-derived adipocytes (n=6). **b,** mRNA analysis of *Fabp4*, *Adipoq*, *Pparg*, *Ppargc1a*, and *Cebpa* in WT and FI3KO SVF-derived adipocytes (n=5). **c,** Lipolysis in WT and FI3KO SVF-derived adipocytes treated with or without insulin (10nM, 4h) (n=5). **d,** Glucose uptake in mouse adipocytes after treatment with varying Poly I:C doses for 2 days (n=8). **e,** Western blot showing phosphorylation of murine IRF3 (S388) in mouse adipocytes after 30 mins of Poly I:C (5 µg/ml) treatment. **f,** Glucose uptake in mouse adipocytes transduced with lentivirus expressing shRNA against *Irf3* or shScr control hairpin in the absence or presence of Poly I:C (5 µg/ml) (n=8). **g,** Glucose uptake in human adipocytes after treatment with varying Poly I:C doses for 2 days (n=8). **h,** Western blot showing phosphorylation of human IRF3 (S396) in human adipocytes after 30 mins of Poly I:C (5 µg/ml) treatment. **i,** Glucose uptake in human adipocytes transduced with lentivirus expressing shRNA against *Irf3* or shScr control hairpin in the absence or presence of Poly I:C (5 µg/ml) (n=8). Statistical significance was assessed by *two-way* ANOVA (**c**, **d** and **g**) or *three-way* ANOVA (**f** and **i**). Data in all panels are expressed as mean ± SEM. *P < 0.05 vs basal, ^#^P < 0.05 vs GFP or shScr.

**Supplemental Fig. 2.**
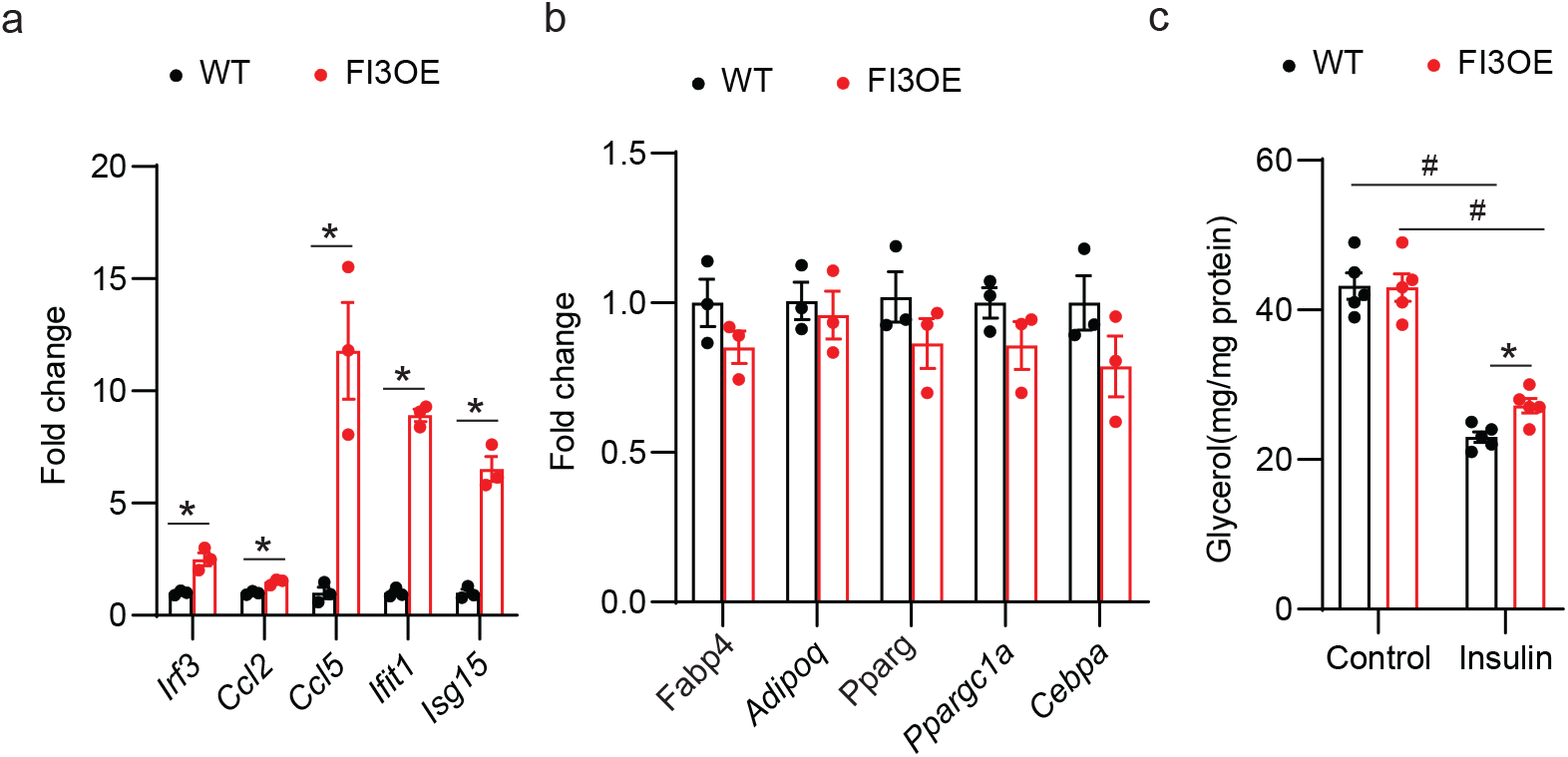
Overexpression of IRF3 does not affect differentiation state in mature adipocytes. **a,** mRNA analysis of *Irf3*, *Ccl5*, *Ifit1*, and *Isg15* in WT and FI3OE SVF-derived adipocytes (n=5). **b,** mRNA analysis of *Fabp4*, *Adipoq*, *Pparg*, *Ppargc1a*, and *Cebpa* in WT and FI3OE SVF-derived adipocytes (n=5). **c,** Lipolysis in WT and FI3OE SVF-derived adipocytes treated with or without insulin (10nM, 4h) (n=5). Statistical significance was assessed by two-tailed Student’s *t-test* (**a**) or *three-way* ANOVA (**c**). Data in all panels are expressed as mean ± SEM. *P < 0.05 vs WT, ^#^P < 0.05 vs control.

**Supplemental Fig. 3.**
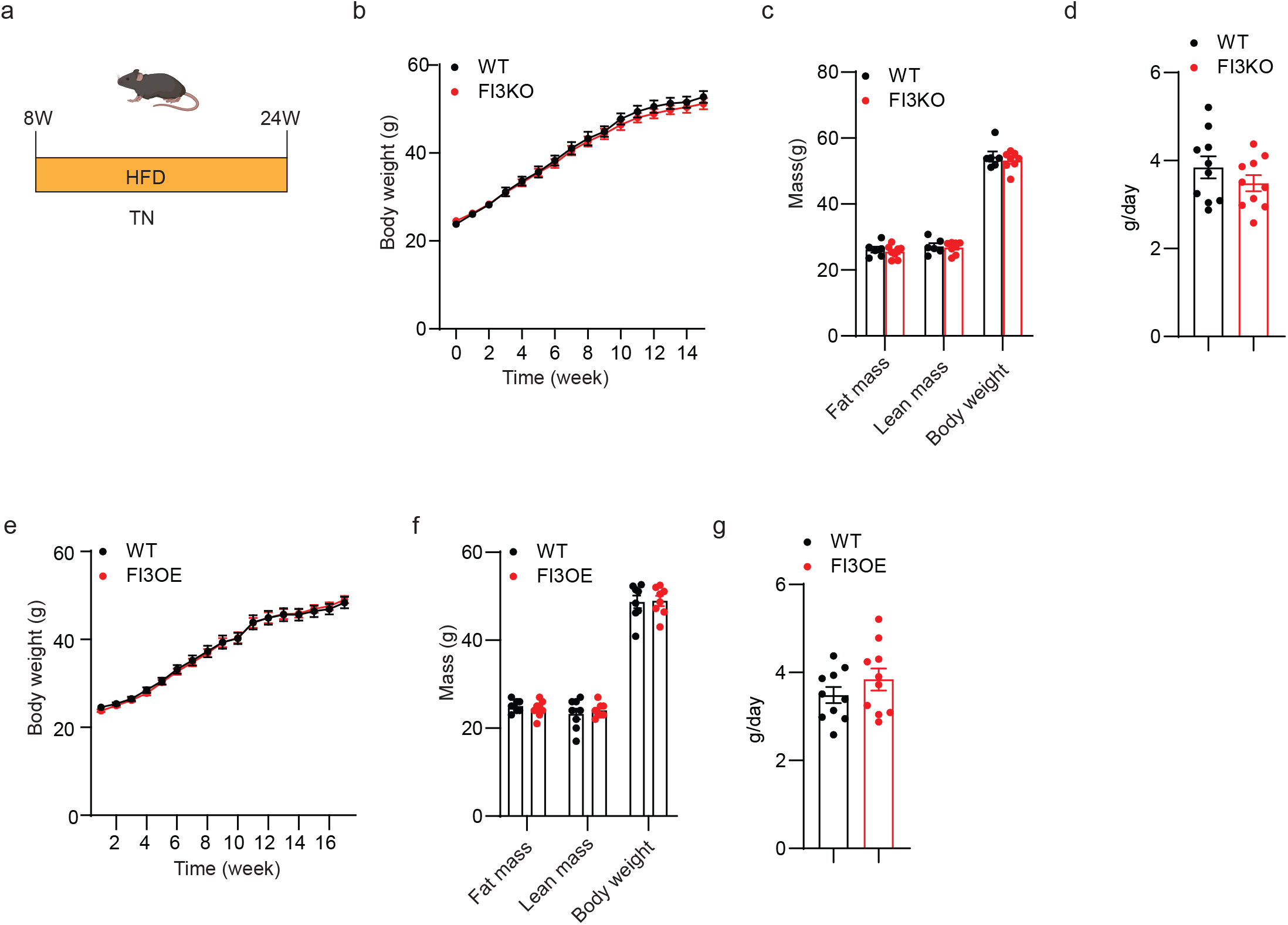
Metabolic phenotype of FI3KO and FI3OE mice on HFD in thermoneutrality. (**a**-**c**) Metabolic analysis of mice as described in Fig. 3a, body weight (**a**), body composition (**b**), and food intake (**c**). (**e**-**g**) Metabolic analysis of mice as described in Fig. 4a, body weight (**e**), body composition (**f**), and food intake (**g**).

**Supplemental Fig. 4.**
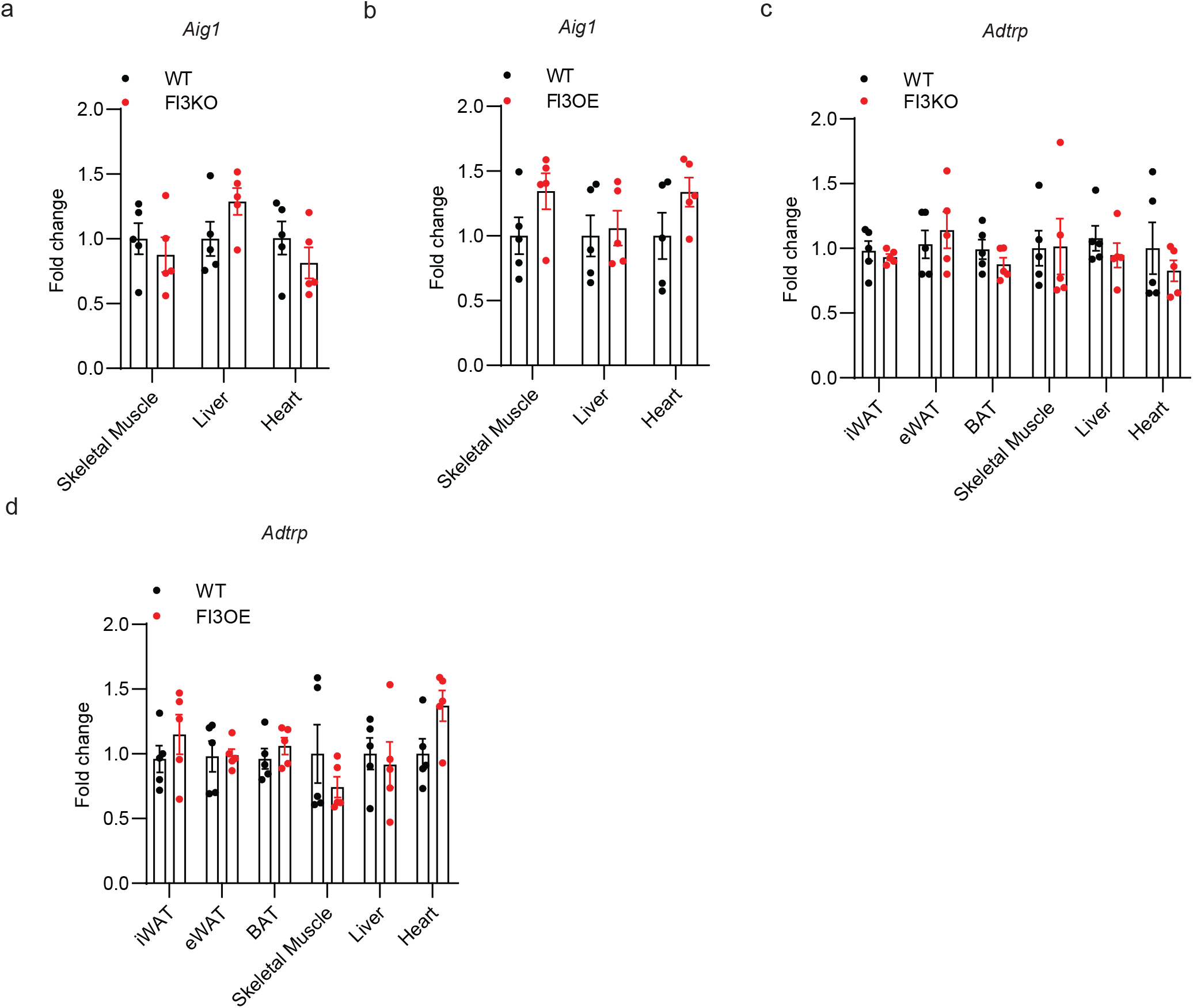
IRF3 regulates AIG1 transcription in adipocytes. **a,** mRNA analysis of *Aig1* in skeletal muscle, liver, and heart of WT and FI3KO mice (n=5). **b,** mRNA analysis of *Aig1* in skeletal muscle, liver, and heart of WT and FI3OE mice (n=5). **c,** mRNA analysis of *Adtrp* in skeletal muscle, liver, and heart of WT and FI3KO mice (n=5). **d,** mRNA analysis of *Adtrp* in skeletal muscle, liver, and heart of WT and FI3OE mice (n=5).

**Supplemental Fig. 5.**
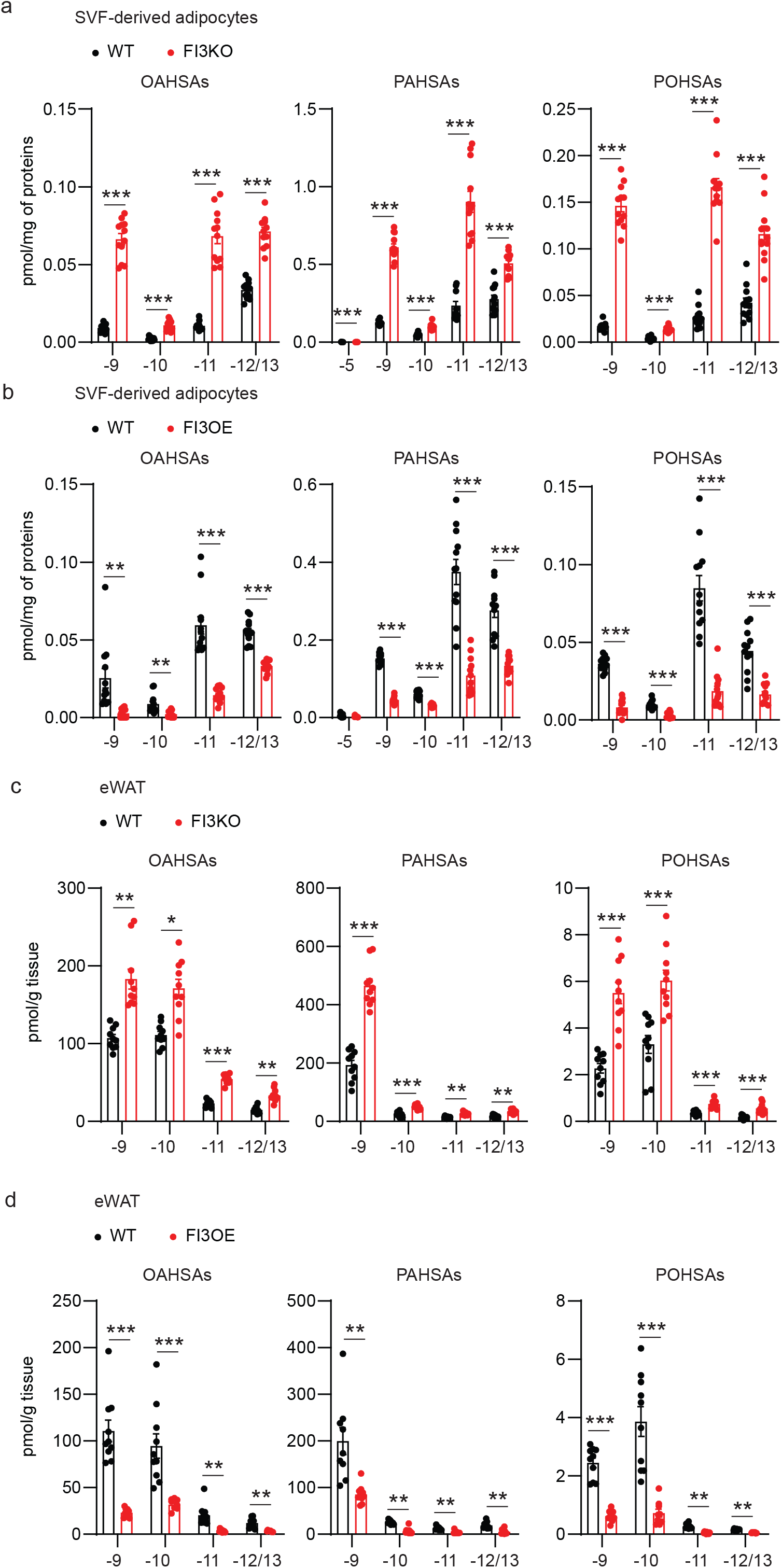
IRF3 decreases FAHFAs in adipocytes. **a,** Quantification of FAHFA isomers in WT and FI3KO SVF-derived adipocytes (n=12). **b,** Quantification of FAHFA isomers in WT and FI3OE SVF-derived adipocytes (n=12). **c,** Quantification of FAHFA isomers in eWAT of WT and FI3KO mice (n=10). **d,** Quantification of FAHFA isomers in eWAT of WT and FI3OE mice (n=10). Statistical significance was assessed by two-tailed Student’s *t-test*. Data in all panels are expressed as mean ± SEM. *P < 0.05 vs WT.

**Supplemental Fig. 6.**
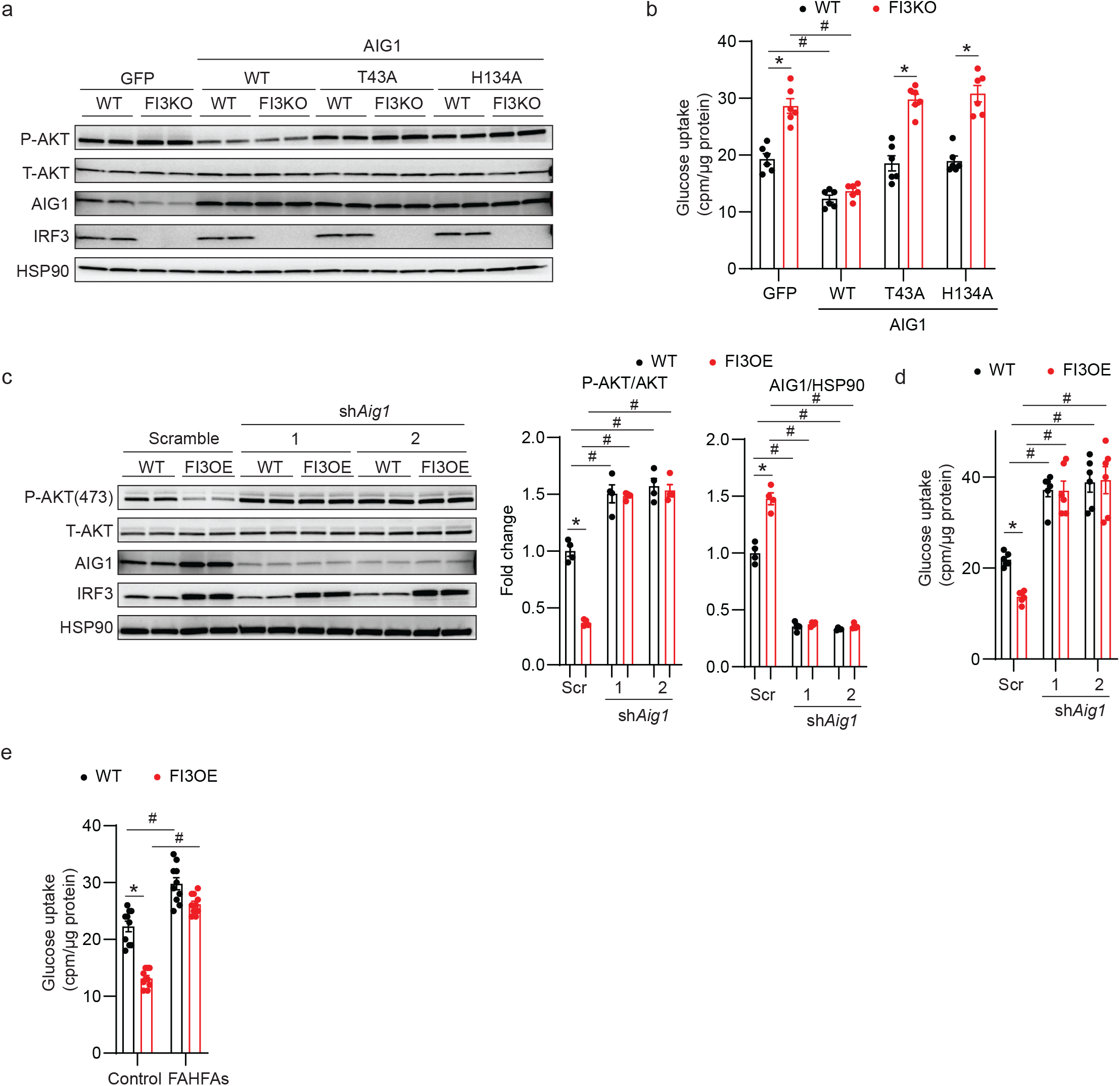
IRF3 promotes insulin resistance through AIG1 in adipocytes. **a,** Western blot of pAKT (S473) and IRF3, all lanes in the presence of insulin, **b,** Insulin-stimulated glucose uptake (n=6) in WT and FI3KO SVF-derived adipocytes infected with GFP, WT AIG1, T43A AIG1, or H134A AIG1 lentivirus. **c,** Western blot of pAKT (S473) and IRF3, **d,** Insulin-stimulated glucose uptake (n=6) in WT and FI3OE SVF-derived adipocytes infected lentivirus expressing shRNA against *Aig1* or shScr control hairpin. **e,** Insulin-stimulated glucose uptake in WT and FI3OE SVF-derived adipocytes treated with control or FAHFAs [9-PAHSA (20µM), 9-POHSA (20µM), and 9-OAHSA (20µM); total FAHFA levels 60μM] for 24h. Statistical significance was assessed by *two-way* ANOVA (**b**-**e**). Data in all panels are expressed as mean ± SEM. *P < 0.05 vs WT, ^#^P < 0.05 vs GFP or shScr.

**Supplemental Fig. 7.**
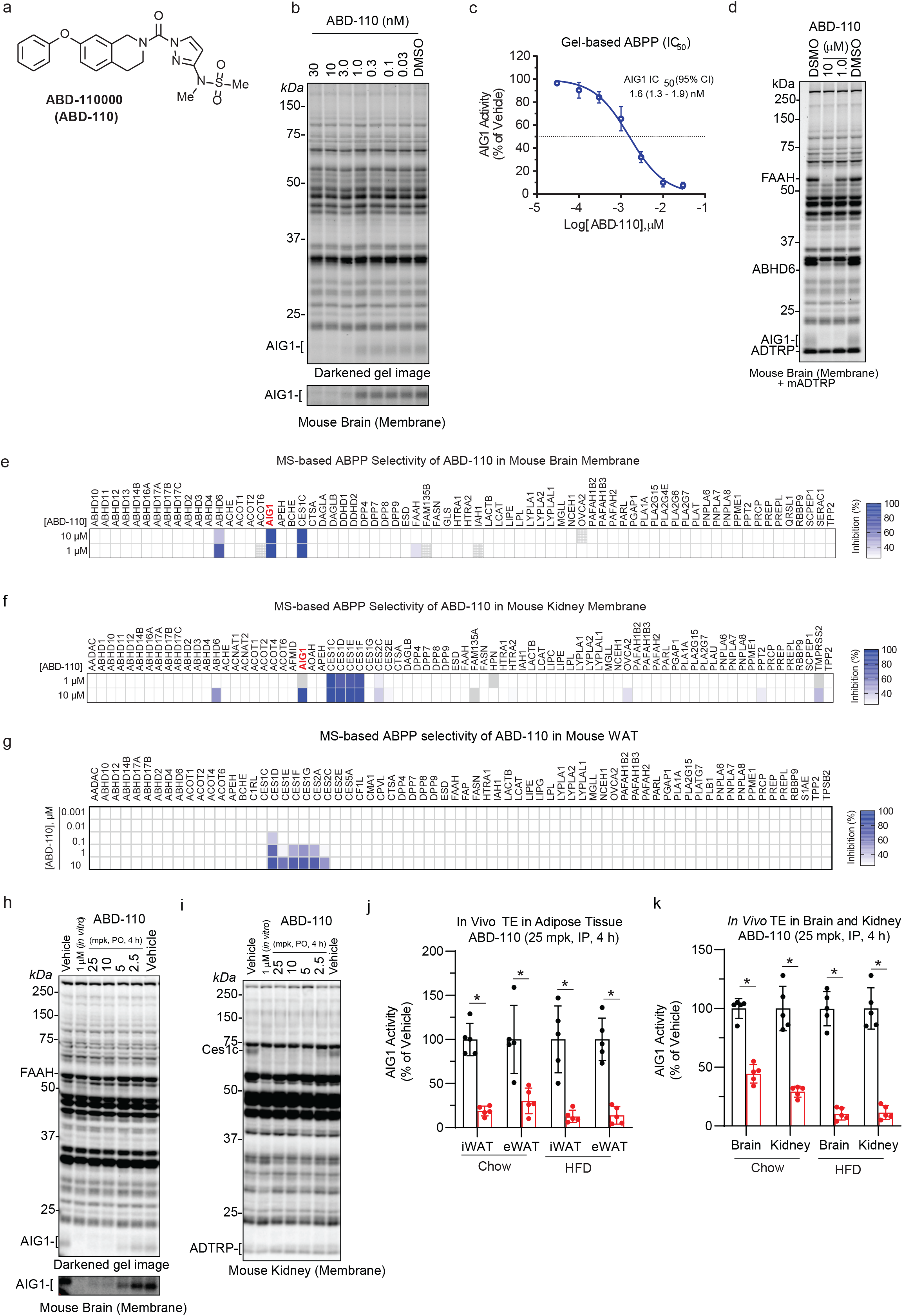
Characterization of the AIG1 inhibitor ABD-110000. **a,** Chemical structure of ABD-110000 (ABD-110). **b,** Competitive gel-based ABPP profile of ABD-110 (0.03–30 nM, 30 min) in mouse brain membrane proteome using FP-Rh. **c,** ABD-110 potency (IC50) for AIG1 determined by competitive gel-based ABPP. Error bars represent SD from three replicates. **d,** Selectivity of ABD-110 (1 & 10 μM, 30 min) against FP-Rh-reactive enzymes in mouse brain membrane proteomes spiked with mADTRP-transfected HEK293T cell proteomes. **e**, **f,** In-depth selectivity profile of ABD-110 (1.0 and 10 μM) across serine/threonine hydrolase enzymes using MS-based ABPP profiles in mouse brain membrane (**e**) and kidney membrane (**f**) proteomes. Data presented represent mean inhibition from three replicates. Gray indicates targets that were not detected in any replicate. **g**, In-depth selectivity profile of ABD-110 (0.001-10 μM, 1 h) across serine hydrolase enzymes using MS-based ABPP profiles in Chow-fed mouse eWAT proteomes. Data presented represent mean inhibition from three replicates. **h, i,** Gel-based ABPP profiles in brain (**h**) and kidney (**i**) membrane proteomes derived from C57Bl/6J mice treated with either vehicle or ABD-110 (2.5 - 25 mpk, PO) for 4 h. Selective AIG1 inhibition is observed in brain whereas kidney profiles, where AIG1 is not detectable, reveal high selectivity across additional peripheral serine hydrolases and ADTRP. **j**, **k,** *In vivo* AIG1 target engagement following ABD-110 administration. Chow or HFD-fed mice were dosed with vehicle or ABD-110 (25 mpk, IP) once daily for 2 weeks. Brain, kidney, eWAT, and iWAT tissues were harvested four hours after the final dose for MS-Based ABPP analysis. AIG1 activity was determined by measuring the degree of activity-dependent AIG1 enrichment relative to the vehicle using parallel reaction monitoring (PRM) to detect and quantify diagnostic AIG1 peptides. Statistical significance was assessed by two-tailed Student’s *t-test*. *P < 0.05 vs vehicle.

**Supplemental Fig. 8.**
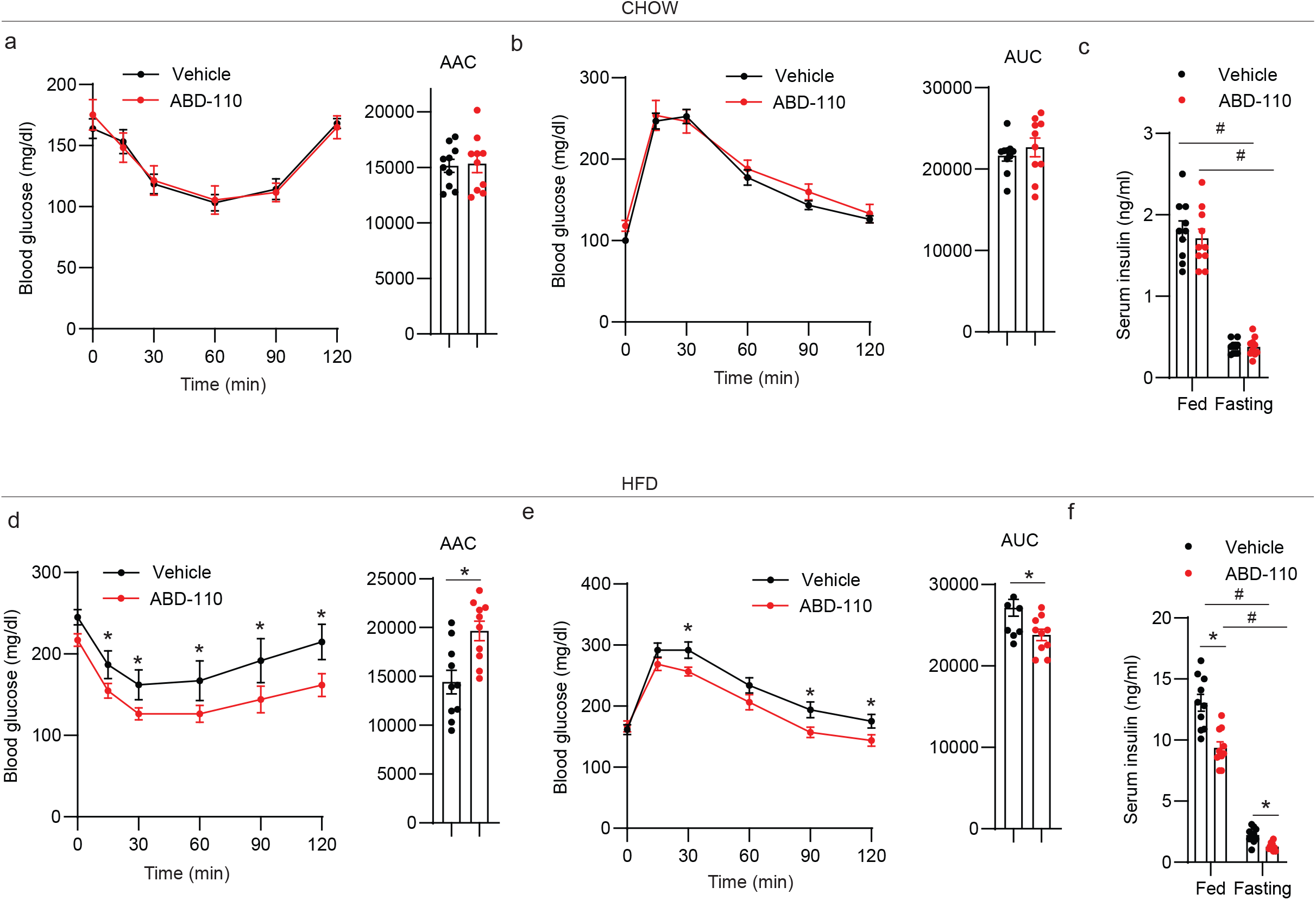
Metabolic effects of ABD-110 on WT Chow and HFD mice. Metabolic analysis of male WT mice (n=10) dosed with vehicle or ABD-110 (25 mg/kg IP) once daily for 2 weeks, including insulin tolerance test (**a**), glucose tolerance test (**b**), *ad lib* fed serum insulin levels (c). Metabolic analysis of male WT mice (n=10) after 16 weeks of HFD feeding dosed with vehicle or ABD-110 (25 mg/kg IP) once daily for 2 weeks, including insulin tolerance test (**d**), glucose tolerance test (**e**), *ad lib* fed serum insulin levels (f). Statistical significance was assessed by *two-way* ANOVA (**c and f**) or two-tailed Student’s *t-test* (**d** and **e**). Data in all panels are expressed as mean ± SEM. *P < 0.05 vs vehicle.

**Supplemental Fig. 9.**
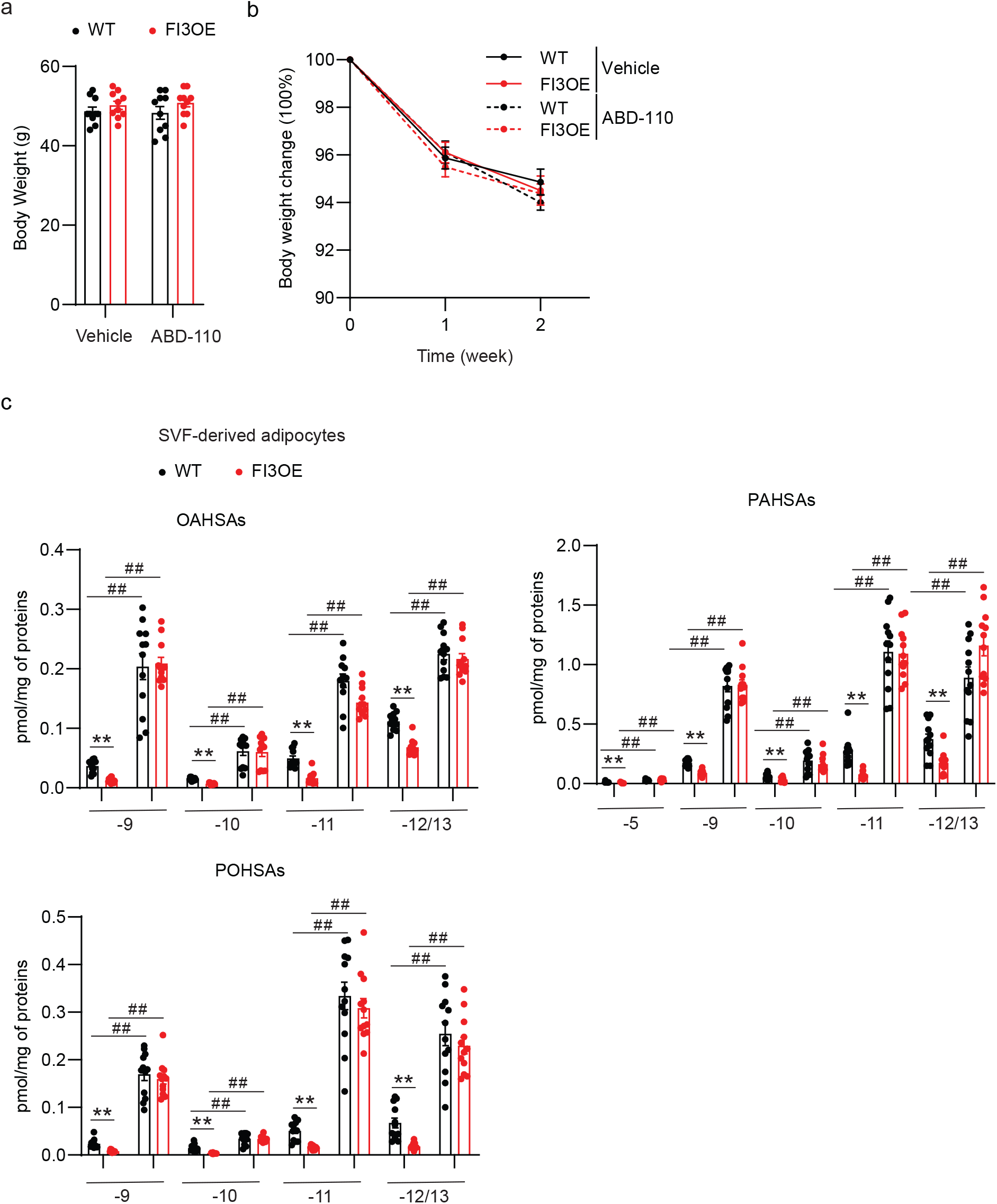
Inhibition of AIG1 by ABD-110 ameliorates HFD-induced insulin resistance in FI3OE mice. **a,** Body weight change of WT and FI3OE mice receiving a daily injection of vehicle or ABD-110 for two weeks (n=7-8). **b,** Body weight of WT and FI3OE mice after receiving a daily injection of vehicle or ABD-110 for two weeks (n=7-8). **c,** Quantification of FAHFA isomers in WT and FI3OE SVF-derived adipocytes treated with or without ABD-110 (1μM, 4h) (n=12). Data in all panels are expressed as mean ± SEM. **P < 0.01 vs WT, ^##^P < 0.01 vs GFP or DMSO.

## References

1 Reilly, S. M. & Saltiel, A. R. Adapting to obesity with adipose tissue inflammation. Nat Rev Endocrinol 13, 633–643 (2017). https://doi.org:10.1038/nrendo.2017.90

2 Weisberg, S. P. et al. CCR2 modulates inflammatory and metabolic effects of high-fat feeding. J Clin Invest 116, 115–124 (2006). https://doi.org:10.1172/JCI24335

3 Oh, D. Y., Morinaga, H., Talukdar, S., Bae, E. J. & Olefsky, J. M. Increased macrophage migration into adipose tissue in obese mice. Diabetes 61, 346–354 (2012). https://doi.org:10.2337/db11-0860

4 Amano, S. U. et al. Local proliferation of macrophages contributes to obesity-associated adipose tissue inflammation. Cell Metab 19, 162–171 (2014). https://doi.org:10.1016/j.cmet.2013.11.017

5 Wu, H. & Ballantyne, C. M. Metabolic Inflammation and Insulin Resistance in Obesity. Circ Res 126, 1549–1564 (2020). https://doi.org:10.1161/CIRCRESAHA.119.315896

6 Nishimoto, S. et al. Obesity-induced DNA released from adipocytes stimulates chronic adipose tissue inflammation and insulin resistance. Sci Adv 2, e1501332 (2016). https://doi.org:10.1126/sciadv.1501332

7 Hersoug, L. G., Moller, P. & Loft, S. Role of microbiota-derived lipopolysaccharide in adipose tissue inflammation, adipocyte size and pyroptosis during obesity. Nutr Res Rev 31, 153–163 (2018). https://doi.org:10.1017/S0954422417000269

8 Kang, S. et al. Identification of nuclear hormone receptor pathways causing insulin resistance by transcriptional and epigenomic analysis. Nat Cell Biol 17, 44–56 (2015). https://doi.org:10.1038/ncb3080

9 Kang, S., Tsai, L. T. & Rosen, E. D. Nuclear Mechanisms of Insulin Resistance. Trends Cell Biol 26, 341–351 (2016). https://doi.org:10.1016/j.tcb.2016.01.002

10 Kitamoto, T., Kuo, T., Okabe, A., Kaneda, A. & Accili, D. An integrative transcriptional logic model of hepatic insulin resistance. Proc Natl Acad Sci U S A 118 (2021). https://doi.org:10.1073/pnas.2102222118

11 James, D. E., Stockli, J. & Birnbaum, M. J. The aetiology and molecular landscape of insulin resistance. Nat Rev Mol Cell Biol 22, 751–771 (2021). https://doi.org:10.1038/s41580-021-00390-6

12 Vallabhapurapu, S. & Karin, M. Regulation and function of NF-kappaB transcription factors in the immune system. Annu Rev Immunol 27, 693–733 (2009). https://doi.org:10.1146/annurev.immunol.021908.132641

13 Cai, D. et al. Local and systemic insulin resistance resulting from hepatic activation of IKK-beta and NF-kappaB. Nat Med 11, 183–190 (2005). https://doi.org:10.1038/nm1166

14 Gao, Z. et al. Serine phosphorylation of insulin receptor substrate 1 by inhibitor kappa B kinase complex. J Biol Chem 277, 48115–48121 (2002). https://doi.org:10.1074/jbc.M209459200

15 Jiao, P. et al. Constitutive activation of IKKbeta in adipose tissue prevents diet-induced obesity in mice. Endocrinology 153, 154–165 (2012). https://doi.org:10.1210/en.2011-1346

16 Gao, Z. et al. P65 inactivation in adipocytes and macrophages attenuates adipose inflammatory response in lean but not in obese mice. Am J Physiol Endocrinol Metab 308, E496–505 (2015). https://doi.org:10.1152/ajpendo.00532.2014

17 Gao, Z. et al. Inactivation of NF-kappaB p50 leads to insulin sensitization in liver through post-translational inhibition of p70S6K. J Biol Chem 284, 18368–18376 (2009). https://doi.org:10.1074/jbc.M109.007260

18 Minegishi, Y. et al. Deletion of nuclear factor-kappaB p50 upregulates fatty acid utilization and contributes to an anti-obesity and high-endurance phenotype in mice. Am J Physiol Endocrinol Metab 309, E523–533 (2015). https://doi.org:10.1152/ajpendo.00071.2015

19 Tamura, T., Yanai, H., Savitsky, D. & Taniguchi, T. The IRF family transcription factors in immunity and oncogenesis. Annu Rev Immunol 26, 535–584 (2008). https://doi.org:10.1146/annurev.immunol.26.021607.090400

20 Eguchi, J. et al. Interferon regulatory factors are transcriptional regulators of adipogenesis. Cell Metab 7, 86–94 (2008). https://doi.org:10.1016/j.cmet.2007.11.002

21 Eguchi, J. et al. Transcriptional control of adipose lipid handling by IRF4. Cell Metab 13, 249–259 (2011). https://doi.org:10.1016/j.cmet.2011.02.005

22 Kong, X. et al. IRF4 is a key thermogenic transcriptional partner of PGC-1alpha. Cell 158, 69–83 (2014). https://doi.org:10.1016/j.cell.2014.04.049

23 Jefferies, C. A. Regulating IRFs in IFN Driven Disease. Front Immunol 10, 325 (2019). https://doi.org:10.3389/fimmu.2019.00325

24 Kumari, M. et al. IRF3 promotes adipose inflammation and insulin resistance and represses browning. J Clin Invest 126, 2839–2854 (2016). https://doi.org:10.1172/JCI86080

25 Yan, S. et al. IRF3 reduces adipose thermogenesis via ISG15-mediated reprogramming of glycolysis. J Clin Invest 131 (2021). https://doi.org:10.1172/JCI144888

26 Patel, S. J. et al. Hepatic IRF3 fuels dysglycemia in obesity through direct regulation of Ppp2r1b. Sci Transl Med 14, eabh3831 (2022). https://doi.org:10.1126/scitranslmed.abh3831

27 McKernan, K., Varghese, M., Patel, R. & Singer, K. Role of TLR4 in the induction of inflammatory changes in adipocytes and macrophages. Adipocyte 9, 212–222 (2020). https://doi.org:10.1080/21623945.2020.1760674

28 Lin, R., Heylbroeck, C., Pitha, P. M. & Hiscott, J. Virus-dependent phosphorylation of the IRF-3 transcription factor regulates nuclear translocation, transactivation potential, and proteasome-mediated degradation. Mol Cell Biol 18, 2986–2996 (1998). https://doi.org:10.1128/MCB.18.5.2986

29 Jiao, Y. et al. Discovering metabolic disease gene interactions by correlated effects on cellular morphology. Mol Metab 24, 108–119 (2019). https://doi.org:10.1016/j.molmet.2019.03.001

30 Doyle, S. et al. IRF3 mediates a TLR3/TLR4-specific antiviral gene program. Immunity 17, 251–263 (2002). https://doi.org:10.1016/s1074-7613(02)00390-4

31 Ganeshan, K. & Chawla, A. Warming the mouse to model human diseases. Nat Rev Endocrinol 13, 458–465 (2017). https://doi.org:10.1038/nrendo.2017.48

32 Yore, M. M. et al. Discovery of a class of endogenous mammalian lipids with anti-diabetic and anti-inflammatory effects. Cell 159, 318–332 (2014). https://doi.org:10.1016/j.cell.2014.09.035

33 Parsons, W. H. et al. AIG1 and ADTRP are atypical integral membrane hydrolases that degrade bioactive FAHFAs. Nat Chem Biol 12, 367–372 (2016). https://doi.org:10.1038/nchembio.2051

34 Erikci Ertunc, M., et al. AIG1 and ADTRP are endogenous hydrolases of fatty acid esters of hydroxy fatty acids (FAHFAs) in mice. J Biol Chem 295, 5891–5905 (2020). https://doi.org:10.1074/jbc.RA119.012145

35 Alexander, J. P. & Cravatt, B. F. The putative endocannabinoid transport blocker LY2183240 is a potent inhibitor of FAAH and several other brain serine hydrolases. J Am Chem Soc 128, 9699–9704 (2006). https://doi.org:10.1021/ja062999h

36 Zvonok, N. et al. Covalent inhibitors of human monoacylglycerol lipase: ligand-assisted characterization of the catalytic site by mass spectrometry and mutational analysis. Chem Biol 15, 854–862 (2008). https://doi.org:10.1016/j.chembiol.2008.06.008

37 Bachovchin, D. A. et al. Superfamily-wide portrait of serine hydrolase inhibition achieved by library-versus-library screening. Proc Natl Acad Sci U S A 107, 20941–20946 (2010). https://doi.org:10.1073/pnas.1011663107

38 Hotamisligil, G. S. et al. IRS-1-mediated inhibition of insulin receptor tyrosine kinase activity in TNF-alpha- and obesity-induced insulin resistance. Science 271, 665–668 (1996). https://doi.org:10.1126/science.271.5249.665

39 Uysal, K. T., Wiesbrock, S. M., Marino, M. W. & Hotamisligil, G. S. Protection from obesity-induced insulin resistance in mice lacking TNF-alpha function. Nature 389, 610–614 (1997). https://doi.org:10.1038/39335

40 Cani, P. D. et al. Metabolic endotoxemia initiates obesity and insulin resistance. Diabetes 56, 1761–1772 (2007). https://doi.org:10.2337/db06-1491

41 Sabio, G. & Davis, R. J. cJun NH2-terminal kinase 1 (JNK1): roles in metabolic regulation of insulin resistance. Trends Biochem Sci 35, 490–496 (2010). https://doi.org:10.1016/j.tibs.2010.04.004

42 Solinas, G. & Becattini, B. JNK at the crossroad of obesity, insulin resistance, and cell stress response. Mol Metab 6, 174–184 (2017). https://doi.org:10.1016/j.molmet.2016.12.001

43 Arkan, M. C. et al. IKK-beta links inflammation to obesity-induced insulin resistance. Nat Med 11, 191–198 (2005). https://doi.org:10.1038/nm1185

44 Conwell, L. S., Trost, S. G., Brown, W. J. & Batch, J. A. Indexes of insulin resistance and secretion in obese children and adolescents: a validation study. Diabetes Care 27, 314–319 (2004). https://doi.org:10.2337/diacare.27.2.314

45 Quinn, C. E., Hamilton, P. K., Lockhart, C. J. & McVeigh, G. E. Thiazolidinediones: effects on insulin resistance and the cardiovascular system. Br J Pharmacol 153, 636–645 (2008). https://doi.org:10.1038/sj.bjp.0707452

46 Abel, E. D. et al. Adipose-selective targeting of the GLUT4 gene impairs insulin action in muscle and liver. Nature 409, 729–733 (2001). https://doi.org:10.1038/35055575

47 Herman, M. A. et al. A novel ChREBP isoform in adipose tissue regulates systemic glucose metabolism. Nature 484, 333–338 (2012). https://doi.org:10.1038/nature10986

48 Vijayakumar, A. et al. Absence of Carbohydrate Response Element Binding Protein in Adipocytes Causes Systemic Insulin Resistance and Impairs Glucose Transport. Cell Rep 21, 1021–1035 (2017). https://doi.org:10.1016/j.celrep.2017.09.091

49 Leonardini, A., Laviola, L., Perrini, S., Natalicchio, A. & Giorgino, F. Cross-Talk between PPARgamma and Insulin Signaling and Modulation of Insulin Sensitivity. PPAR Res 2009, 818945 (2009). https://doi.org:10.1155/2009/818945

50 Syed, I. et al. Palmitic Acid Hydroxystearic Acids Activate GPR40, Which Is Involved in Their Beneficial Effects on Glucose Homeostasis. Cell Metab 27, 419–427 e414 (2018). https://doi.org:10.1016/j.cmet.2018.01.001

51 Lee, J. et al. Branched Fatty Acid Esters of Hydroxy Fatty Acids (FAHFAs) Protect against Colitis by Regulating Gut Innate and Adaptive Immune Responses. J Biol Chem 291, 22207–22217 (2016). https://doi.org:10.1074/jbc.M115.703835

52 Patel, R. et al. ATGL is a biosynthetic enzyme for fatty acid esters of hydroxy fatty acids. Nature 606, 968–975 (2022). https://doi.org:10.1038/s41586-022-04787-x

53 Kolar, M. J. et al. Branched Fatty Acid Esters of Hydroxy Fatty Acids Are Preferred Substrates of the MODY8 Protein Carboxyl Ester Lipase. Biochemistry 55, 4636–4641 (2016). https://doi.org:10.1021/acs.biochem.6b00565

54 Emont, M. P. et al. A single-cell atlas of human and mouse white adipose tissue. Nature 603, 926–933 (2022). https://doi.org:10.1038/s41586-022-04518-2

55 Paluchova, V. et al. Lipokine 5-PAHSA Is Regulated by Adipose Triglyceride Lipase and Primes Adipocytes for De Novo Lipogenesis in Mice. Diabetes 69, 300–312 (2020). https://doi.org:10.2337/db19-0494

56 Aryal, P. et al. Distinct biological activities of isomers from several families of branched fatty acid esters of hydroxy fatty acids (FAHFAs). J Lipid Res 62, 100108 (2021). https://doi.org:10.1016/j.jlr.2021.100108

57 Jimenez, V. et al. FGF21 gene therapy as treatment for obesity and insulin resistance. EMBO Mol Med 10 (2018). https://doi.org:10.15252/emmm.201708791

58 Kir, S. et al. Tumour-derived PTH-related protein triggers adipose tissue browning and cancer cachexia. Nature 513, 100–104 (2014). https://doi.org:10.1038/nature13528

59 Chen, S., Zhou, Y., Chen, Y. & Gu, J. fastp: an ultra-fast all-in-one FASTQ preprocessor. Bioinformatics 34, i884–i890 (2018). https://doi.org:10.1093/bioinformatics/bty560

60 Patro, R., Duggal, G., Love, M. I., Irizarry, R. A. & Kingsford, C. Salmon provides fast and bias-aware quantification of transcript expression. Nat Methods 14, 417–419 (2017). https://doi.org:10.1038/nmeth.4197

61 Soneson, C., Love, M. I. & Robinson, M. D. Differential analyses for RNA-seq: transcript-level estimates improve gene-level inferences. F1000Res 4, 1521 (2015). https://doi.org:10.12688/f1000research.7563.2

62 O’Neill, B. T. et al. Differential Role of Insulin/IGF-1 Receptor Signaling in Muscle Growth and Glucose Homeostasis. Cell Rep 11, 1220–1235 (2015). https://doi.org:10.1016/j.celrep.2015.04.037

63 Bligh, E. G. & Dyer, W. J. A rapid method of total lipid extraction and purification. Can J Biochem Physiol 37, 911–917 (1959). https://doi.org:10.1139/o59-099

64 Zhang, T. et al. A LC-MS-based workflow for measurement of branched fatty acid esters of hydroxy fatty acids. Nat Protoc 11, 747–763 (2016). https://doi.org:10.1038/nprot.2016.040

65 Liu, Y., Patricelli, M. P. & Cravatt, B. F. Activity-based protein profiling: the serine hydrolases. Proc Natl Acad Sci U S A 96, 14694–14699 (1999). https://doi.org:10.1073/pnas.96.26.14694

66 Inloes, J. M. et al. The hereditary spastic paraplegia-related enzyme DDHD2 is a principal brain triglyceride lipase. Proc Natl Acad Sci U S A 111, 14924–14929 (2014). https://doi.org:10.1073/pnas.1413706111

67 Zecha, J. et al. TMT Labeling for the Masses: A Robust and Cost-efficient, In-solution Labeling Approach. Mol Cell Proteomics 18, 1468–1478 (2019). https://doi.org:10.1074/mcp.TIR119.001385

68 Gao, J. et al. CIMAGE2.0: An Expanded Tool for Quantitative Analysis of Activity-Based Protein Profiling (ABPP) Data. J Proteome Res 20, 4893–4900 (2021). https://doi.org:10.1021/acs.jproteome.1c00455

69 Hulstaert, N., et al. ThermoRawFileParser: Modular, Scalable, and Cross-Platform RAW File Conversion. J Proteome Res 19, 537–542 (2020). https://doi.org:10.1021/acs.jproteome.9b00328

70 Eng, J. K., Jahan, T. A. & Hoopmann, M. R. Comet: an open-source MS/MS sequence database search tool. Proteomics 13, 22–24 (2013). https://doi.org:10.1002/pmic.201200439

71 UniProt, C. UniProt: the Universal Protein Knowledgebase in 2023. Nucleic Acids Res 51, D523–D531 (2023). https://doi.org:10.1093/nar/gkac1052

72 Tyanova, S., Temu, T. & Cox, J. The MaxQuant computational platform for mass spectrometry-based shotgun proteomics. Nat Protoc 11, 2301–2319 (2016). https://doi.org:10.1038/nprot.2016.136

73 Rost, H. L. et al. OpenMS: a flexible open-source software platform for mass spectrometry data analysis. Nat Methods 13, 741–748 (2016). https://doi.org:10.1038/nmeth.3959

74 The, M., MacCoss, M. J., Noble, W. S. & Kall, L. Fast and Accurate Protein False Discovery Rates on Large-Scale Proteomics Data Sets with Percolator 3.0. J Am Soc Mass Spectrom 27, 1719–1727 (2016). https://doi.org:10.1007/s13361-016-1460-7

